# LuxUS: DNA Methylation Analysis Using Generalized Linear Mixed Model with Spatial Correlation

**DOI:** 10.1101/536722

**Authors:** Viivi Halla-aho, Harri Lähdesmäki

## Abstract

**Motivation:** DNA methylation is an important epigenetic modification, which has multiple functions. DNA methylation and its connections to diseases have been extensively studied in recent years. It is known that DNA methylation levels of neighboring cytosines are correlated and that differential DNA methylation typically occurs rather as regions instead of individual cytosine level.

**Results:** We have developed a generalized linear mixed model, LuxUS, that makes use of the correlation between neighboring cytosines to facilitate analysis of differential methylation. LuxUS implements a likelihood model for bisulfite sequencing data that accounts for experimental variation in underlying biochemistry. LuxUS can model both binary and continuous covariates, and mixed model formulation enables including replicate and cytosine random effects. Spatial correlation is included to the model through a cytosine random effect correlation structure. We show with simulation experiments that by utilizing the spatial correlation we gain more power to the statistical testing of differential DNA methylation. Results with real bisulfite sequencing data set show that LuxUS is able to detect biologically significant differentially methylated cytosines.

**Availability:** The tool is available at https://github.com/hallav/LuxUS.

**Supplementary information:** Supplementary data are available at *bioRxiv*.

## 1 Introduction

DNA methylation is a widely studied epigenetic modification, which is involved in gene regulation. Aberrant methylation states have been shown to be connected with diseases and cancer. One popular method for quantifying DNA methylation levels is bisulfite sequencing (BS-seq) and its variants. Along with the development of sequencing techniques, several computational tools have been proposed for analysis of differential methylation from BS-seq data. These methods aim to model the underlying methylation proportions and perform statistical tests to assess the statistical significance of the effect of a covariate of interest on the methylation level.

Spatial correlation of cytosines’ methylation states is a widely known phenomenon (Eckhardt *et al.*, 2006) and methylation state is generally thought to vary rather as regions instead of individual cytosine level. Consequently, many DNA methylation analysis tools have attempted to capture the phenomenon with different model structures and computational techniques. For example, the popular beta-binomial regression-based tool RADMeth by Dolzhenko and Smith (2014) outputs log-likelihood ratio test p-values for each cytosine separately and then uses weighted Z-test to combine the p-value of a single site with the p-values of its neighbors. RADMeth also includes a feature for merging neighboring differentially methylated cytosines into differentially methylated regions (DMRs). Similarly, Wen *et al.* (2016) first use beta-binomial regression model to calculate p-values for individual CpG sites and then combine them using Getis-Ord statistic. BSmooth tool by Hansen *et al.* (2012), included in bsseq package, is based on local-likelihood smoothing, which can compensate for low-coverage data and then uses signal-to-noise statistic similar to t-test for identifying differentially methylated regions. bsseq package also includes implementation of Fisher’s exact test. DSS tool includes two versions of a beta-binomial model with hypothesis testing with Wald test, one with two-group comparison (Feng *et al.*, 2014) and another with a general experimental design (Park *et al.*, 2016). metilene tool by Jühling *et al.* (2016) utilizes binary segmentation algorithm with two-dimensional Kolmogorov-Smirnov test for finding differentially methylated regions. Recently published dmrseq tool (Korthauer *et al.*, 2018) combines smoothing and covariance structure with weighted least squares regression. Mayo *et al.* (2014) proposed a kernel based method, M^3^D, where statistical testing of putative DMRs is done through maximum mean discrepancy values. Rackham *et al.* (2017) take a different approach with their tool ABBA, which uses a latent Gaussian model.

Another approach without spatial correlation, LuxGLM, was proposed by Äijo *et al.* (2016) and it enables testing for differential methylation for individual cytosines while taking into account different methylcytosine species and experimental design through generalized linear model part. LuxGLM can also utilize spike-ins, such as the commonly used (unmethylated) lambda phage genome, to model and estimate the experimental parameters at the same time with the actual model parameters. In Äijo *et al.* (2016) the LuxGLM tool was shown to be accurate and perform on par or even better than other recent methods in detecting differential methylation.

Here we propose LuxUS, a method where the methylation proportions are modeled with a generalized linear mixed model (GLMM) that can take into account both binary and continuous variables and random effects. Instead of analysing each cytosine separately, we propose to analyse all cytosines in a moderately sized genomic window at a time. The spatial correlation of the neighboring cytosines’ methylation states is included in the model through a covariance structure of the cytosine random effect. To allow individual variation, we also introduce a replicate random effect. After estimating the model parameters, the user can view summary statistics of the posterior distributions of all linear and mixed effect coefficients, allowing for comprehensive evaluation of their significance and proportions.

## 2 Methods

Here we present the LuxUS model, which consists of two parts: an observation model and generalized linear mixed model. Observation model attempts to model the bisulfite sequencing count data generation. The linear model part is used for estimating the methylation proportion parameter for the data generation process. The plate diagram for the whole LuxUS model is presented in Fig. 1. We use Bayesian approach and probabilistic programming language Stan (Carpenter *et al.*, 2017) for implementing the model and estimating the unknown model parameters. Hypothesis testing of the significance of explanatory variables is done using Bayes factors. To choose the genomic windows for the LuxUS analysis we also implemented a simple preprocessing step. The workflow of a bisulfite sequencing data analysis with LuxUS is presented in diagram Supplementary Fig. S3.

**Figure 1:**
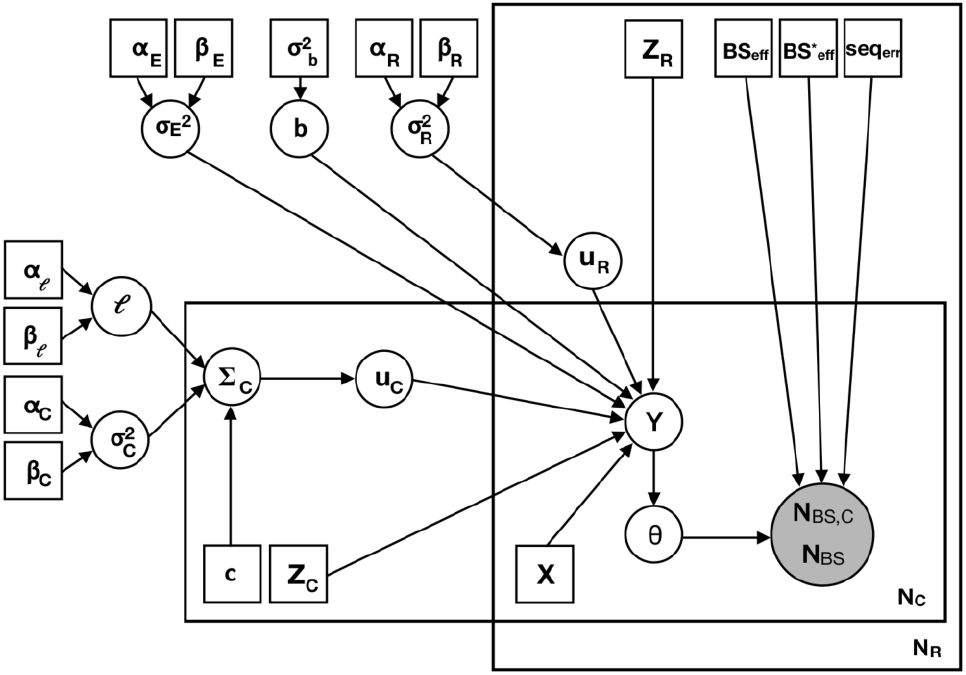
Plate diagram of the LuxUS model. The grey and white circles represent observed and latent variables respectively. Rectangles represent fixed hyperparameters, design matrices or other type of input data.

### 2.1 An observation model for bisulfite sequencing data

We present the LuxUS method for a single genomic window and drop the index denoting a particular window position to simplify notation. The observation model is the same as the one presented in Äijo *et al.* (2016), which we first review here. From bisulfite sequencing data we can retrieve, for each cytosine, the total number of reads overlapping the corresponding cytosine in each sample, **N**_BS_, out of which **N**_BS,C_ observations were cytosines. This is demostrated in Supplementary Fig. S2. Both **N**_BS_ and **N**_BS,C_ are vectors of length *N*_*R*_ · *N*_*C*_, where *N*_*R*_ is the number of samples and *N*_*C*_ is the number of cytosines in the genomic window of interest. In bisulfite sequencing the DNA goes first through a bisulfite conversion treatment, which converts unmethylated cytosines to urasiles with probability BS_Eff_. The methylated cytosines stay unconverted with probability 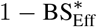. After the bisulfite treatment, DNA is sequenced and the urasiles are observed as thymines (T) and cytosines are observed as cytosines (C). The probability of sequencing error, i.e. observing urasile as C or cytosine as T, is seq_Err_. The experimental parameters seq_Err,*i*_, BS_Eff,*i*_ and 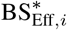 are used in the model to describe the probabilities of sequencing error, bisulfite conversion and incorrect bisulfite conversion, respectively, for the sample corresponding to the *i*th observation. The complete probability tree for observing a C or T in a BS-seq experiment is presented in Suppl. Fig. S1. The probability of observing *N*_BS,C,*i*_ many cytosine reads out of total read count *N*_BS,*i*_ is modelled with binomial distribution with success probability parameter *p*_BS,C,*i*_

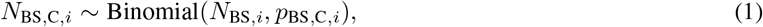

where *i* = 1, …, *N*_*C*_ · *N*_*R*_ and where the elements of 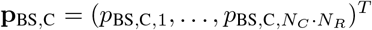 are calculated using the experimental parameters described above

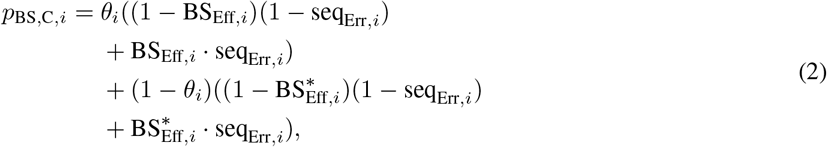

following the probability tree diagram in Suppl. Fig. S1. The *θ*_*i*_ denotes the *i*th element of methylation proportion vector ***θ***, which is modelled with a generalized linear mixed effect model as described in Section 2.2.

It is often thought that samples with low bisulfite conversion efficiency should be excluded from analyses, as they are considered unreliable. But as the experimental parameters can be taken into account with LuxUS, one does not have to leave out such samples from the analysis as long as the conversion efficiencies can be reliably estimated.

### 2.2 A generalized mixed model with spatial correlation

As stated above, the methylation proportions ***θ*** are estimated using a generalized linear mixed model, which includes fixed effect covariates and cytosine and replicate random effects. The number of fixed effect covariates, counting in the possible intercept term, is *N*_*P*_. Here we assume that the effects of the covariates in design matrix **X** are fixed, but each of the replicates and cytosines have their own random intercept terms. A column vector 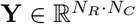 can be expressed as a sum of fixed and random effects

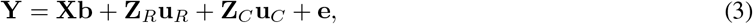

where design matrix **X** of shape (*N*_*C*_ · *N*_*R*_) × *N*_*P*_ and fixed effect coefficient vector **b** of length *N*_*P*_ form the fixed effect term, whereas replicate random effect design matrix **Z**_*R*_ of shape (*N*_*C*_ · *N*_*R*_) × *N*_*R*_ and replicate random effects vector **u**_*R*_ of length *N*_*R*_ form the replicate random effect term and cytosine random effect design matrix **Z**_*C*_ of shape (*N*_*C*_ · *N*_*R*_) × *N*_*C*_ and cytosine random effects vector **u**_*C*_ of length *N*_*C*_ form the cytosine random effect term. The last term, vector **e** of length *N*_*R*_ · *N*_*C*_, is the noise term of the model. The rows of **Y** and design matrices *X*, **Z**_*R*_ and **Z**_*R*_ should all be ordered with the same principle, e.g. by first listing the *N*_*R*_ replicates of the first cytosine, then the replicates of the second cytosine, etc.

Finally, the covariate and random effects are connected to the methylation proportion vector 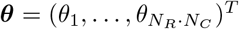 through the sigmoid link function

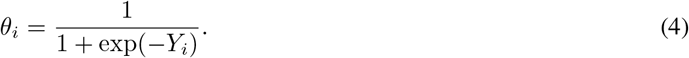

The methylation proportions ***θ*** are used in the calculation of *p*_BS,C,*i*_ as described in Eq. 2.

For the fixed effect, the design matrix **X** contains the experimental design. The *N*_*R*_ × *N*_*P*_ design matrix **D** for one cytosine is used as a block matrix in **X** which is the design matrix of the experiment and applies to the whole genomic window. The rows of the matrix **D** correspond to the replicates in the experiment and the columns correspond to the covariates. The matrix **D** is repeated *N*_*C*_ times in the following way to form **X**

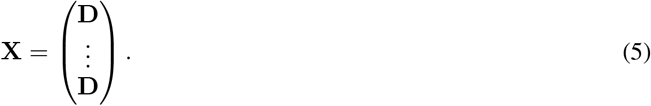

The fixed effect coefficients vector **b** has a normal prior

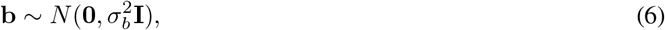

where the prior variance 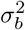 should be set to high enough value to enable sufficiently high variation for the fixed effect coefficients.

The random effect terms for replicates and cytosines are expressed as vectors **u**_*R*_ and **u**_*C*_ with normal priors

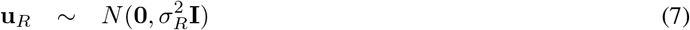

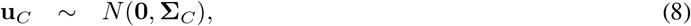

which are multiplied with random effect design matrices **Z**_*R*_ and **Z**_*C*_ respectively in Eq. 3. **Z**_*C*_ and **Z**_*R*_ indicate from which cytosine and replicate each observation is coming from. Random term **u**_*R*_ has a diagonal covariance matrix with variance 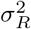 as the diagonal term, whereas **u**_*C*_ has a covariance matrix **Σ**_*C*_ which includes the spatial correlation of the model. The covariance matrix **Σ**_*C*_ is defined as

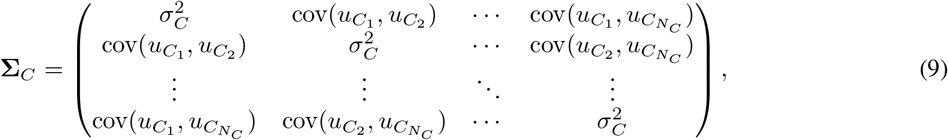

where the covariance terms between the elements of **u**_*C*_ are defined as

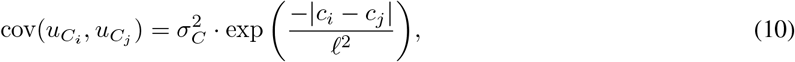

where *i* = 1, …, *N*_*C*_, *j* = 1, …, *N*_*C*_, and the genomic coordinates of the cytosines are stored in vector 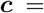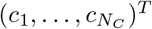. The cytosines in the genomic window are restricted to be located in the same chromosome, as otherwise it would not be possible to calculate genomic distance between the cytosines. The length-scale parameter, *ℓ*, is set a gamma prior

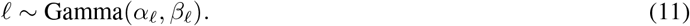

Hyperparameters are set to *α*_*ℓ*_ = 38 and *β*_*ℓ*_ = 1 as they result in covariance that is similar to the methylation correlation plots shown in (Eckhardt *et al.*, 2006; Song *et al.*, 2017). To demonstrate the shape of the covariance term the term 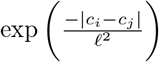 has been plotted as a function of the distance *c*_*i*_ − *c*_*j*_ in Suppl. Fig. S4.

The random effect variance parameters 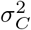 and 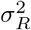 have gamma priors

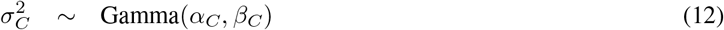

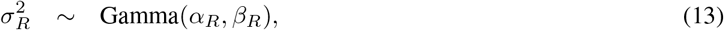

where *α*_*R*_, *β*_*R*_, *α*_*C*_ and *β*_*C*_ are set to make the prior distribution match our prior information about the cytosine and replicate random effects. The residual noise term **e** is defined similarly

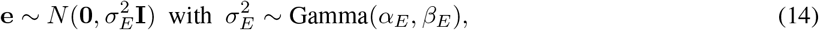

where *α*_*E*_ and *β*_*E*_ are the shape and rate parameters of gamma distribution, respectively. We have also implemented an alternative version of the model where inverse-gamma distributions are used instead of the gamma priors for the parameters 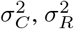 and 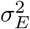.

With the above definitions for the fixed and random effect, the distribution of **Y** can be expressed as

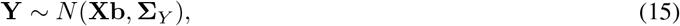

where the elements of the covariance matrix **Σ**_*Y*_ are defined as

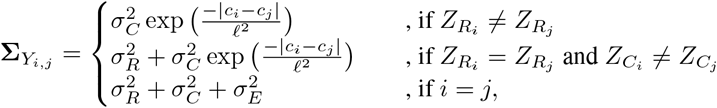

where *i* = 1, …, *N*_*R*_ · *N*_*C*_ and *j* = 1, …, *N*_*R*_ · *N*_*C*_ and 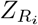 and 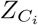 denote the *i*th rows of random effect design matrices **Z**_*R*_ and **Z**_*C*_ respectively.

### 2.3 Model estimation

For further inference based on the model, we have to estimate the unknown parameters. The variables to be estimated are fixed effect coefficients **b**, length-scale parameter *ℓ*, replicate random effect variance 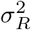, cytosine random effect variance 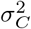, noise term variance 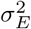, replicate random effect **u**_*R*_, cytosine random effect **u**_*C*_ and linear predictor **Y**, from which methylation proportion ***θ*** can be calculated as described in Eq. 4.

The model was implemented with probabilistic programming language Stan (Carpenter *et al.*, 2017) and the model parameters are estimated using the no-U-turn sampling algorithm, which is a locally adaptive version of Hamiltonian Monte Carlo (HMC) sampling as implemented in Stan. The PyStan version (Stan Development Team, 2017) of the software was used for the HMC sampling. Stan also has a built-in automatic differentiation variational inference (ADVI) feature (Kucukelbir *et al.*, 2015), which was also tested for estimating the model parameters. The mean-field algorithm, which is the default option in ADVI, was used. A more detailed description of model estimation algorithms can be found from Suppl. file Section S1.1. As variational inference approaches are often faster than Markov chain Monte Carlo (MCMC) methods, it could be a more favourable model estimation method especially when analysing large reduced representation bisulfite sequencing (RRBS-seq) or whole genome bisulfite sequencing (WGBS-seq) data sets. For running ADVI we utilize CmdStan (Stan Development Team, 2018), which is the command line interface to Stan.

### 2.4 Testing for differential methylation

After estimating the model parameters as described above, differential methylation can be tested. With the models described above it is possible to perform two types of tests. The type 1 test has null hypothesis *H*_0_ : **b**_*i*_ = 0 and alternative hypothesis *H*_1_ : **b**_*i*_ ≠ 0, i.e. the statistical significance of the covariate *i* is tested. The type 2 test has null hypothesis *H*_0_ : **b**_*i*_ − **b**_*j*_ = 0 and alternative hypothesis *H*_1_ : **b**_*i*_ − **b**_*j*_ ≠ 0, i.e. the difference between the effects of covariates *i* and *j* is tested for statistical significance. The testing is done using Bayes factors. As calculating the exact values of Bayes factors is often infeasible, instead the Savage-Dickey density ratio estimate of the Bayes factor is used. For the type 1 test the Savage-Dickey estimate is

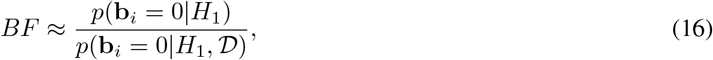

where we denote data with 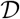. The Savage-Dickey estimator for the type 2 test is defined similarly. The numerator of the Savage-Dickey density ratio is analytically solvable as we use normal prior for **b**. As for the denominator, it can be approximated from the posterior samples for **b**_*i*_ by evaluating a kernel density estimate of the posterior at origin. The Gaussian kernel density estimation function from statistical functions module from the SciPy package (Virtanen *et al.*, 2020) is used to calculate the kernel density estimates. Scott’s rule is used as the bandwidth parameter for the kernel density estimation. The Bayes factor is calculated for the whole window being analyzed, as the linear model coefficients **b** are shared among all cytosines.

### 2.5 Preanalysis step for determining and filtering the genomic windows

To prepare a BS-seq experiment data set for LuxUS analysis, we have provided a script which also includes a simple preanalysis filtering method. The genomic windows, for which the analysis is performed, can be determined using a fixed window size (in terms of number of nucleotides) or number of cytosines in a window. The prefiltering step is computationally efficient and can be used to filter away regions which e.g. do not contain sufficient number of reads or are unlikely to exhibit differential DNA methylation. A coverage limit can be used to filter out cytosines with low coverages. As estimating the model for Bayes factor calculation can be computationally burdensome in genome-wide studies, we have implemented an F-test for testing the significance of a variable of interest using the logit transformed sample means (calculated over all the cytosines in the window) of the methylation states. The p-value limit for accepting a window for further analysis with the LuxUS model can be set as desired by the user. Each genomic window can then be analysed further separately, enabling parallelisation.

## 3 Results

To demonstrate the performance of the LuxUS model and to compare it to other tools, we applied LuxUS to real and simulated BS-seq data sets. First, the results from analysis of colon cancer WGBS-seq data set for LuxUS and RADMeth were compared. The performance of LuxUS, RADMeth, M^3^D, bsseq, DSS and metilene were compared on simulated data sets. For LuxUS, both HMC and ADVI approaches for model fitting were tested. To show the advantage of using spatial correlation structure, each cytosine in the simulated data sets was also analysed separately.

### 3.1 Analysis of colon cancer WGBS-seq data

The model was tested using a WGBS-seq data set by Hansen (2018), consisting of matched human colon and colon cancer samples. The processed sequencing data was provided for two chromosomes, 21 and 22. First, the genomic windows for LuxUS analysis were determined using the preanalysis method. The details of the preanalysis and setting the priors can be found from Suppl. file Section S2.1. Then Stan was run with HMC sampling to fit the LuxUS model parameters and finally Bayes factors were calculated. The sampling was done with four chains with 1500 samples in each, out of which half were discarded as burn-in. As a result 1422 genomic windows were discovered with Bayes factor ≥ 3. These genomic windows covered 21127 cytosines in total. Kass and Raftery (1995) proposed that Bayes factor values from 3 to 20 would indicate positive evidence against the null hypothesis, while values from 20 to 150 indicate strong evidence and values higher than 150 indicate very strong evidence. These limits can be used as approximate guidelines, as LuxUS computes the Savage-Dickey estimates of the Bayes factors instead of the exact Bayes factors.

In comparison, we performed RADMeth analysis on the same data set and discovered 137142 cytosines with adjusted, FDR-corrected p-value ≤ 0.05. The overlap of the significant cytosines from LuxUS and RADMeth approaches is 15008 cytosines, i.e., about three-fourths of the differentially methylated cytosines found by LuxUS are also found by RADMeth. The results for a genomic region in chromosome 21 are shown in Fig. 2, which demonstrates that both LuxUS and RADMeth produce similar results, but LuxUS is more conservative. For example, near the end of the genomic region in Fig. 2 the difference in average methylation state between the cancer and normal colon cells decreases, which RADMeth fails to detect and produces p-values smaller than 0.05. Whereas the said region was filtered from the analysis by LuxUS already in the preanalysis phase. The boxplots of the computation times (on one standard computing node in a cluster) for each number of cytosines in a genomic window for LuxUS are shown in Suppl. Fig. S5. The total computation time on a single core for LuxUS for the 9945 genomic windows was 1623.80 hours. Note that LuxUS supports full parallelization across all genomic windows, although the computation time here is reported for a single computing core. Running RADMeth for the whole dataset took 31 minutes and 33 seconds.

**Figure 2:**
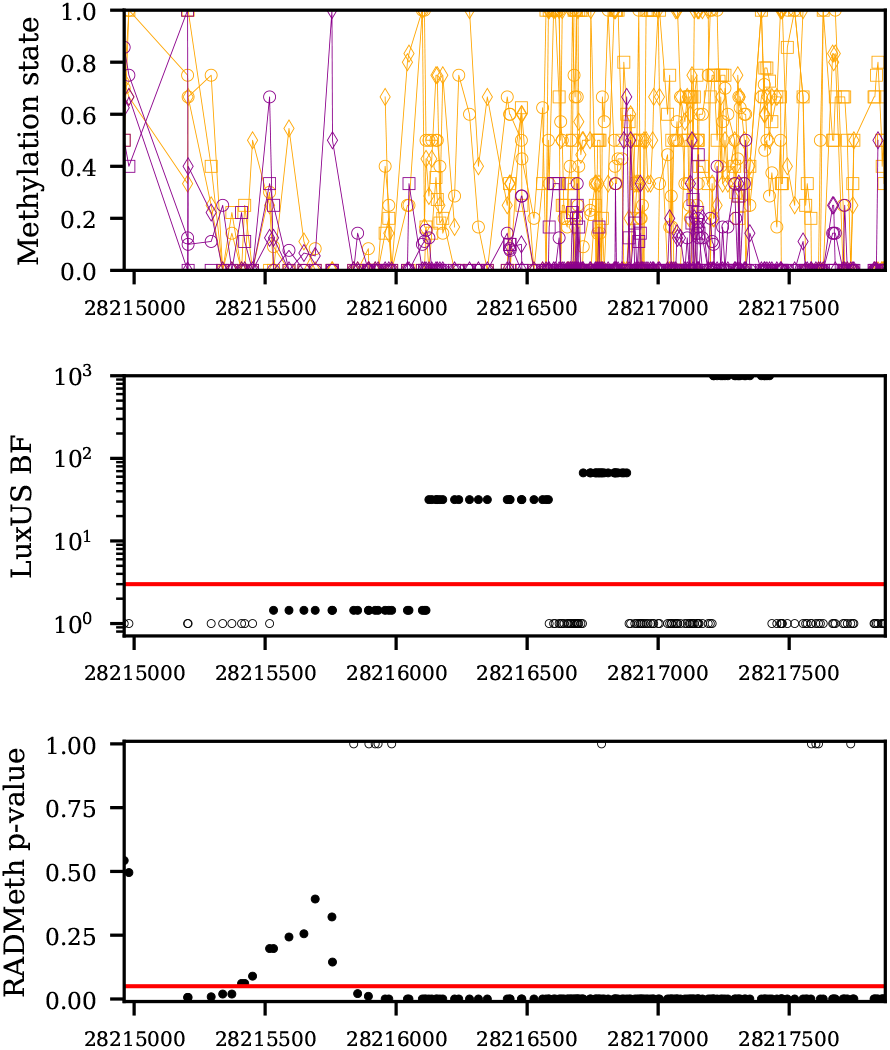
The top panel shows the methylation states for a chosen genomic region in chromosome 21 (from 28214962 bp to 28217867 bp) for the six samples in the colon cancer data set. The colon cancer sample methylation states have been plotted in orange and the normal colon samples in purple. The experiment was paired, and the cancer and colon samples from the same person have been plotted with the same marker symbols. The middle panel shows the LuxUS Bayes factor values with log scaling. The Bayes factor values have been truncated to 1000. The cytosines which were filtered from further analysis in the preanalysis phase have been plotted with value 0 with empty markers. The horizontal red line corresponds to the Bayes factor value 3. The bottom panel shows the RADMeth p-values for the same genomic region. The horizontal red line corresponds to the adjusted p-value of 0.05. The cytosines for which RADMeth produced no p-value are plotted with p-value of 1 with empty markers.

To investigate the biological significance of the found differentially methylated cytosines, we used the GREAT tool (McLean *et al.*, 2010) for gene set enrichment analysis (online version 3.0.0 with the default parameter settings). The lists of significantly differentially methylated cytosines found by LuxUS and RADMeth were given to the tool as inputs and the unfiltered list of CpG sites for the chromosomes 21 and 22 was used as background regions file. The results from both methods showed enrichment in cancer related terms in Disease Ontology and Molecular Signatures Database (MSigDB) Perturbation ontology, which indicates that the methods were able to discover truly differentially methylated cytosines. The lists of the enriched terms for LuxUS and RADMeth can be found from Suppl. Fig S6-S10. Suppl. file Section S2.3 describes how LuxUS preanalysis step was validated using the colon cancer data set.

In Suppl. file Section S2.4 the estimated variance parameters were utilized for setting simulation parameters. To further investigate the effect of the variance term priors, we applied LuxUS to another BS-seq data set by Hascher *et al.* (2014). We also experimented using different 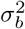 values to see how they affect the calculated Bayes factors and consequently the performance of the method with this data set. These experiments are described in detail in Suppl. file Section S3.

### 3.2 Performance comparisons on simulated BS-seq data

Simulated BS-seq data was generated for performance comparison purposes. The simulations were made using the LuxUS model, which takes into account various mechanisms that affect the real bisulfite sequencing process, such as experimental parameters. First, 100 sets of experimental parameters and cytosine locations were generated. The number of cytosines was 10 and their locations were randomly chosen from range [1, 1000]. The choice of cytosine frequency is explained in detail in Suppl. file Section S2.4. Two BS-seq data sets were then simulated for each of these 100 sets: one with differential methylation and one with no differential methylation between the case and control samples. These 200 simulated genomic regions form a simulated data set with 50% proportion of differentially methylated regions. The values for the fixed effect coefficient means, ***μ***_*b*_ = (*μ*_*B*,0_, *μ*_*B*,1_), for the cases with differential methylation were (−1.4, 1), (−1.4, 2.3), (1.4, −1) and (1.4, −2.3). These values correspond to approximate values of 0.2, 0.5, −0.2 and 0.5 of Δ*θ*, the difference in methylation proportions between the case and control groups, respectively. The variance for the fixed effect coefficients, 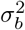, was set to 0.25 for the simulations. The variances for the cytosine and replicate random effects and the noise term, 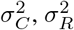 and 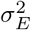, were set to the means of the gamma distributions which were fitted to the posterior samples from the colon cancer WGBS-seq data analysis. Detailed description of this can be found from Suppl. file Section S2.4. The coverage (i.e., the number of reads) as well as the number of replicates were varied as 6, 12 and 24. The priors for the variance terms 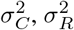 and 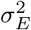 were set using the results from the real data analysis and 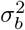 is set to 15 for the model estimation. The detailed description of the generation of the simulated data can be found from Suppl. file Section S4.1.

The model parameters were then estimated using the ADVI and HMC methods in Stan. The HMC sampler was run with four chains and the number of posterior samples for each chain was 1000, out of which half were discarded as burn-in. When running ADVI, 2000 samples were generated from the approximate posterior distribution to match the number of samples retrieved from HMC sampler. The number of gradient samples was 10 and number of samples used for evidence lower bound (ELBO) estimation was 200. These parameter values were chosen based on the results presented in Malonzo *et al.* (2018), where the precision and computation times of ADVI were compared to HMC for the LuxGLM model. After estimating the model parameters, the Bayes factors for the type 1 test were calculated. For comparison, the analysis was run for each cytosine separately to see the difference with the model with added spatial correlation. For this purpose the cytosine random effect was removed from the model, as there was only one cytosine being analysed at a time. The resulting model corresponds to the LuxGLM method by Äijo *et al.* (2016) except that GLM is changed to GLMM by adding the replicate random effect into the model.

To compare our method to other published tools, we ran the analysis also with RADMeth, M^3^D, dmrseq, DSS, metilene and bsseq. RADMeth was run with default parameters and the p-value adjusting was done for each simulated data set separately. Presumably because of the beta-binomial nature of the RADMeth model, there were some cases where the methylation states of both the case and control groups were exactly 0 or 1 for which RADMeth failed to calculate p-values. The result for these cytosines were removed before ROC and AUROC calculations. dmrseq was run first with default parameters and then with a higher maximum number of permutations and lower cutoff value for the single CpG coefficient, but the AUROC values for the method remained very low (possibly for some technical incompatibility reasons) and were thus left out from the result table. M^3^D was run with default parameters. For M^3^D all the simulated data sets (with one simulation setting) were given to M^3^D at the same time as separate testing regions, each with different dummy chromosome name. For the ROC and AUROC calculations the unadjusted p-values were used, as they gave slightly better result than the FDR-adjusted p-values even though the FDR-correction should not change the ranking of the p-values. metilene, bsseq and DSS were all run to compare the difference between case and control groups for each simulated genomic window separately. metilene was run with de-novo DMR finding mode, with maximum distance between CpGs in a DMR set to 1000 and minimum number of CpGs in a DMR set to one. Minimum mean methylation difference for calling DMRs was set to zero. Additionally, we ran metilene with pre-defined regions mode with the same settings as for the default de-novo DMR finding mode, using the start and end coordinates of each simulated genomic window for defining the regions for which the statistical testing is performed. From now on we will refer to the de-novo DMR finding mode as metilene, while the pre-defined regions mode is referred to as metilene mode 2. With bsseq we first ran the smoothing step with minimum number of cytosines in a smoothing window set to one. Then Fisher’s exact test was run with bsseq. The t-test feature in bsseq was not utilized for comparison, as the tool does not include p-value calculation for the t-test statistics. We ran DSS tool with no smoothing and with smoothing, using two different smoothing window spans, 500bp and 1000bp. The approach with 1000bp smoothing window span showed the best performance. The most interesting tools to compare LuxUS with are perhaps RADMeth and DSS, which alike LuxUS allow including additional covariates into the analysis.

The area under receiver operating characteristic curve (AUROC) values for the different approaches are presented in Table 1-2. The ADVI version of LuxUS has a tendency of outputting very high or infinite valued Bayes factors for easier cases. To avoid problems of comparing infinite values to each other in the ROC calculation, the infinite values have been replaced with the greatest non-infinite Bayes factor value with small constant added. For the cytosines for which RADMeth failed to calculate p-values, the values were removed. If the DMRs found by metilene with de-novo DMR finding mode did not cover all the cytosines in a simulated genomic window, the not covered cytosines were given p-value 1 for ROC calculation. The AUROC values in Table 1, presenting results for simulation setting where expected methylation level difference between case and control groups was Δ*θ* ≈ −0.2, show that metilene with pre-defined regions mode slightly outperforms LuxUS in most of the cases. While in Table 2, presenting results for simulation setting where Δ*θ* ≈ −0.5, for the most of the cases the LuxUS model gives the highest AUROC values. The difference between the results from running LuxUS for a genomic window and for each cytosine separately is considerable. For these simulation settings, using variational inference for estimating the model parameters results in nearly as good results as using Hamiltonian Monte Carlo sampling, while also being significantly faster as can be seen from Suppl. Table S1 in which the comparison of mean computation times for the HMC and ADVI approaches is presented. RADMeth, M^3^D, metilene with de-novo DMR finding mode and bsseq show better performance than LuxUS run separately for each cytosine, but they do not achieve the same accuracy as LuxUS and metilene mode 2, with one exception where M^3^D tops all the other methods. DSS performs well, but it has the best AUROC values in only four of the simulation cases shown in the tables. While for LuxUS, RADMeth, metilene, DSS and bsseq performance gets systematically better when the replicate or read count is increases, the performance of M^3^D fluctuates. LuxUS and metilene mode 2 seem to do equally well in this simulation experiment. However, unlike LuxUS metilene is not able to take into account additional covariates which is often desired in real data analysis. The AUROC values and ROC curves for the simulation settings with different *μ*_*b*_ can be found from Suppl. Table S2-S3 and Suppl. Fig S20-S23 respectively. Suppl. file Section S4.7 demonstrates how LuxUS can estimate the underlying methylation proportions. To demostrate that LuxUS performs well even with a smaller, more realistic DMR proportion, we also performed comparisons with a data set consisting of 1000 simulated regions out of which 50 (5%) were DMRs. The results are presented in the form of precision-recall curves and average precision tables in Suppl. file Sections S4.5 and S4.6. The results are fairly similar to the ones with 50% DMR proportion, as LuxUS HMC, metilene and DSS are performing approximately equally well when the methylation state difference between case and control samples is small. When the difference is bigger, LuxUS HMC shows the best performance. RADMeth and M^3^D perform relatively well, while bsseq and LuxUS run separately for each cytosine have the lowest AUROC values. However, this time LuxUS ADVI was not able to reach the same AUROC value levels as LuxUS HMC and its precision-recall curves show unexpected behaviour, which might be because of its tendency of returning very high BF approximation values.

**Table 1:** AUROC values for LuxUS with HMC and ADVI, LuxUS for separate cytosines, RADMeth, M^3^D, metilene, DSS and bsseq for simulated data set with ***μ***_*b*_ = (1.4, −1), corresponding to Δ*θ* ≈ −0.2. The highest AUROC value is bolded for each simulation scenario. *N*_BS_ denotes the number of sequencing reads overlapping a cytosine and *N*_*R*_ denotes the number of samples.

| $N_{BS}$ | $N_R$ | LuxUS<br>HMC | LuxUS<br>ADVI | LuxUS<br>sep. | RAD-Meth | M<sup>3</sup>D | metilene | metilene<br>mode 2 | DSS | bsseq |
| --- | --- | --- | --- | --- | --- | --- | --- | --- | --- | --- |
| 6 | 6 | 0.734 | 0.738 | 0.522 | 0.607 | 0.719 | 0.631 | **0.767** | 0.738 | 0.615 |
| 6 | 12 | 0.814 | 0.783 | 0.606 | 0.712 | 0.703 | 0.658 | **0.818** | 0.801 | 0.643 |
| 6 | 24 | **0.923** | 0.909 | 0.687 | 0.819 | 0.682 | 0.759 | 0.917 | 0.904 | 0.711 |
| 12 | 6 | 0.687 | 0.696 | 0.545 | 0.632 | 0.640 | 0.590 | **0.698** | 0.690 | 0.590 |
| 12 | 12 | 0.822 | 0.803 | 0.625 | 0.756 | 0.701 | 0.678 | **0.827** | 0.812 | 0.654 |
| 12 | 24 | 0.927 | 0.899 | 0.721 | 0.867 | 0.755 | 0.781 | **0.928** | 0.924 | 0.754 |
| 24 | 6 | 0.667 | 0.668 | 0.541 | 0.612 | **0.703** | 0.635 | 0.686 | 0.680 | 0.597 |
| 24 | 12 | 0.840 | 0.835 | 0.637 | 0.765 | 0.777 | 0.735 | 0.850 | **0.852** | 0.712 |
| 24 | 24 | 0.933 | 0.886 | 0.762 | 0.890 | 0.732 | 0.830 | **0.934** | 0.916 | 0.773 |

**Table 2:**
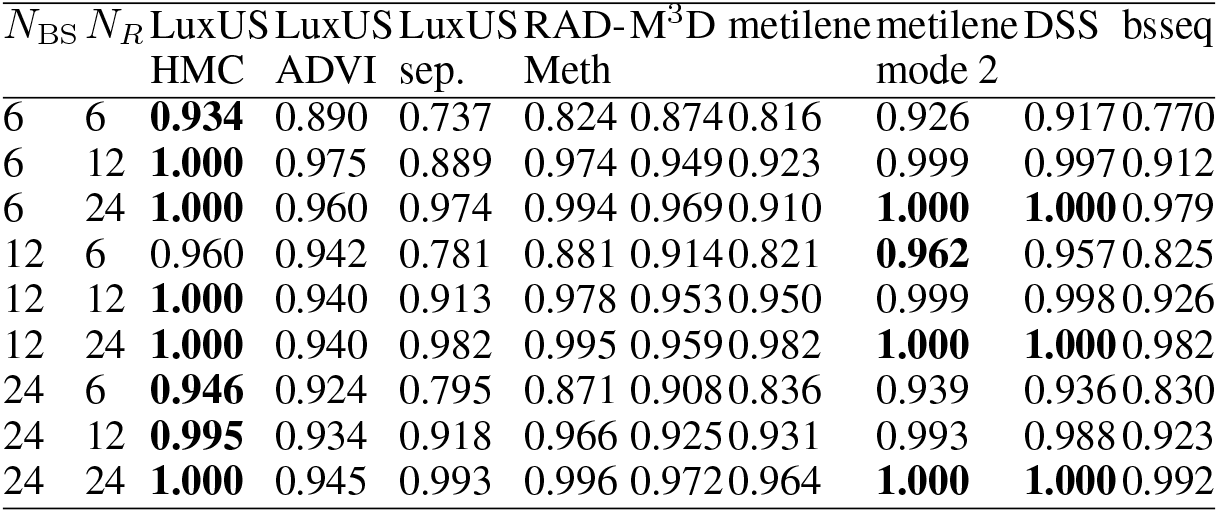
AUROC values for LuxUS with HMC and ADVI, LuxUS for separate cytosines, RADMeth, M^3^D, DSS, metilene and bsseq for simulated data set with *μ*_*B*_ = (1.4, −2.3), corresponding to Δ*θ* ≈ −0.5. The highest AUROC value is bolded for each simulation scenario. *N*_BS_ denotes the number of sequencing reads overlapping a cytosine and *N*_*R*_ denotes the number of samples.

To better demonstrate the concept of adding spatial correlation into the model, we simulated data with different numbers of cytosines in a window of 1000 basepairs and calculated the AUROC values for each case. The coefficient mean ***μ***_*b*_ was set to (1.4, −1) and the number of reads and replicates was 12. Using this simulated data, the LuxUS model was estimated with HMC algorithm and Bayes factors were calculated. The results of this experiment can be seen in Fig. 3, which shows that analysing multiple cytosines at a time gives more statistical power and results in higher AUROC values. The turning point for this simulation setting seems to be between 6 and 8 cytosines, after which adding more cytosines does not seem to add more power at least in this specific case. As the number of cytosines in the analysis increases, so does the number of parameters in the model and the size of the covariance matrices. This increases computation time during model estimation and can also lead to estimation inaccuracies or convergence issues during sampling. The increase in computation time for the colon cancer data set is shown in Suppl. Fig. S5, where the computation times for different number of cytosines being analysed at a time are presented. The results shown in Fig. 3 indicate that increasing the number of cytosines in the analysis after some point does not give more power to the analysis, and it is not advisable in the sense of computational efficiency either.

**Figure 3:**
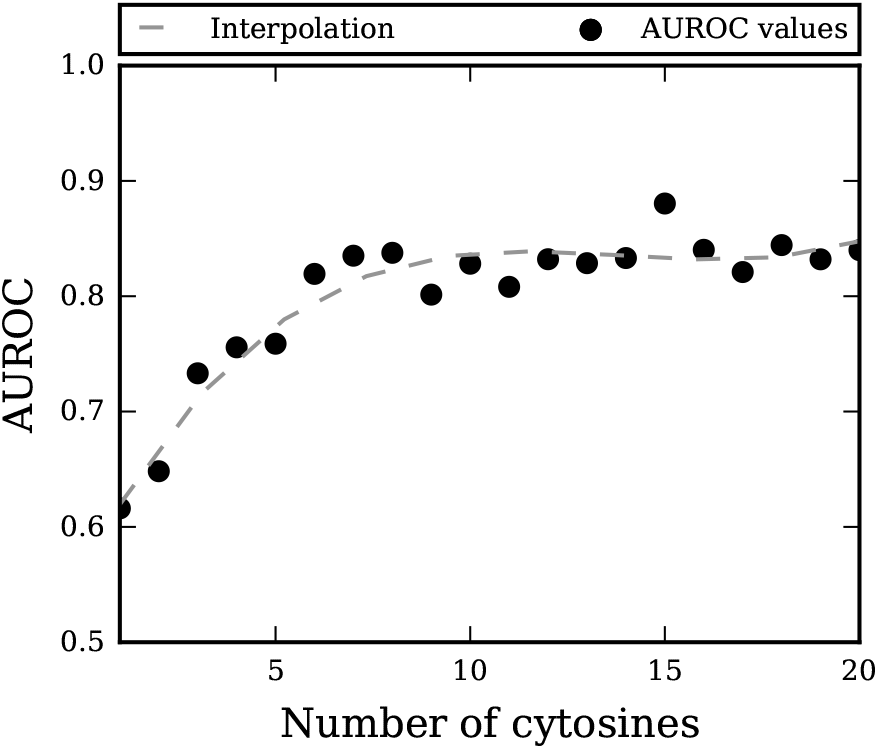
The AUROC value for LuxUS as function of number of cytosines in a 1000 bp window for simulated data and its interpolation with third degree polynomial. Each AUROC value is estimated using 400 data sets, including 200 data sets with differential methylation and 200 data sets with no differential methylation.

LuxUS allows a general experimental design to be used in the analysis, and to present the advantages of this feature we generated a simulated data set with two confounding covariates. The design matrix consists of an intercept term, a binary covariate distinguishing cases from controls, a confounding binary covariate and a confounding continuous covariate, with corresponding coefficients *μ*_*b*_ = (1.4, −1, 2, −3) used in generating the data. Details of the data generation can be found from Suppl. file Section S4.1. The comparisons were conducted for both 50% and 5% DMR proportions. The AUROC values for each method for the comparison with 50% DMR proportion (200 simulated genomic regions in total) are presented in Tab. 3. Corresponding ROC figures can be found from Suppl. file Section S4.8. Based on the AUROC values, LuxUS HMC performs the best, while LuxUS ADVI and DSS tools also do well in the comparison. The continuous covariate was transformed into a binary covariate so that it could be utilized by RADMeth, but its AUROC values are lower than for LuxUS and DSS, which both allow continuous covariates. metilene mode 2 is performing approximately equally well as RADMeth. Unlike with the simple experimental design, the AUROC values for metilene mode 2 are considerably lower than for LuxUS and DSS for this simulation setting where the confounding coefficients have significant effects, muddling the difference between cases and controls. The default mode of metilene, bsseq, M^3^D and LuxUS run separately for each cytosine have the lowest AUROC values, which is expected as metilene, bsseq and M^3^D tools cannot take confounding covariates into account and bsseq and LuxUS (sep.) cannot utilize spatial correlation. The precision-recall curves and average precision tables for the 5% DMR proportion setting (1000 simulated genomic regions in total) are presented in Suppl. file Sections S4.9 and S4.10. Comparing the average precision values, LuxUS HMC shows again the best performance. In this setting, metilene mode 2 seems to be doing slightly better than RADMeth. The average precisions for LuxUS ADVI are again notably lower than for LuxUS HMC in this setting where the DMR proportion is small.

**Table 3:**
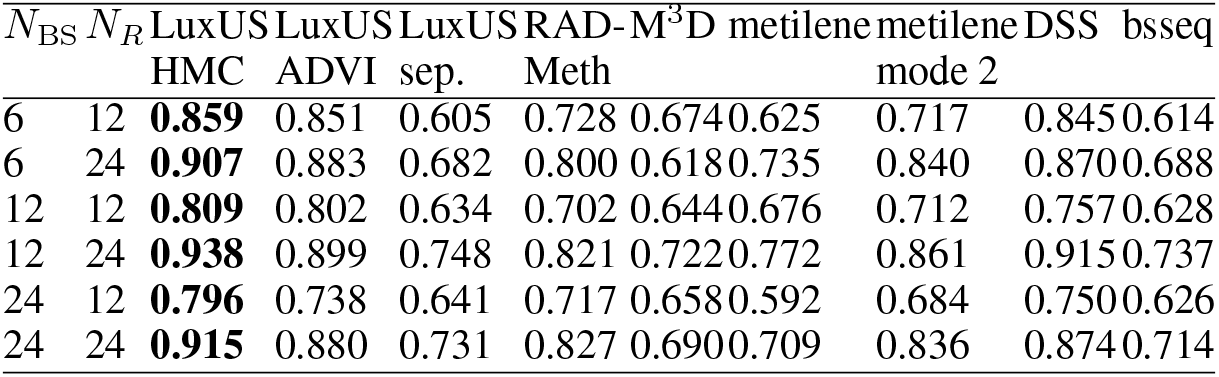
AUROC values for LuxUS with HMC and ADVI, LuxUS for separate cytosines, RADMeth, M^3^D, DSS, metilene and bsseq for simulated data set with confounding covariates. The highest AUROC value is bolded for each simulation scenario. *N*_BS_ denotes the number of sequencing reads overlapping a cytosine and *N*_*R*_ denotes the number of samples.

### 3.3 Performance comparisons on simulated differentially methylated regions based on real BS-seq data

To make further comparisons of the performance of the tools with a model-independent data set, we applied them on simulated data by Hebestreit *et al.* (2013). The simulation was based on RRBS-seq data of 12 control samples. Six of the controls were turned to cases by adding 10000 simulated differentially methylated regions to randomly chosen CpG islands. The size of the DMRs and the methylation difference between the cases and controls was varied. For this comparison the CpG islands containing a DMR were divided into DMR and non-DMR sets. For both sets the LuxUS preanalysis step was run to divide them into genomic windows with LuxUS preanalysis method. The details of the preanalysis and data filtering can be found from Suppl. file Section S5.1. 3000 windows with minimum number of two cytosines were chosen randomly from both DMR and non-DMR groups. LuxUS, RADMeth, M^3^D, bsseq, metilene and DSS analyses were conducted on these 6000 windows. The experimental design consisted of an intercept term and a case-control indicator variable for LuxUS and RADMeth. With M^3^D, bsseq, DSS and metilene the difference between the case and control groups was tested. For RADMeth, the p-value adjustment step was performed for all the 6000 genomic windows together. LuxUS was run with HMC sampler, using four chains with 1500 samples in each out of which half were discarded as burn-in. With DSS tool, three different smoothing window spans (500bp, 1000bp and 2000bp) were tested along with approach with no smoothing. The DSS approach that yielded best AUROC value was smoothing with 500bp window span and was chosen to be presented here. metilene tool was run with de-novo DMR distance between two CpGs in a DMR was increased to 2000bp. bsseq tool was used to run smoothing and Fisher’s finding and pre-defined regions modes with the same settings as described in Section 3.2, that the maximum allowed exact test, similarly as described in Section 3.2. For metilene, DSS and bsseq tools the picked DMR and non-DMR windows were all analysed together after sorting them based on their genomic locations.

As the differentially methylated regions were known for this simulated data set, ROC computation could be performed. Fig. 4 shows that all methods perform approximately equally well. LuxUS (HMC) has the highest AUROC value of 0.992, LuxUS with ADVI model fitting comes in second with AUROC value of 0.980 and RADMeth is third with AUROC value 0.968. metilene, M^3^D and DSS with 500bp smoothing window span all performed well with AUROC values 0.957, 0.938 and 0.937 respectively, while bsseq has the lowest AUROC value of 0.909. For this data set, LuxUS with ADVI produced a great number of very high and infinite valued Bayes factors. The infinite valued BFs were again replaced with the highest non-infinite BF value with a small constant added for the ROC calculation.

**Figure 4:**
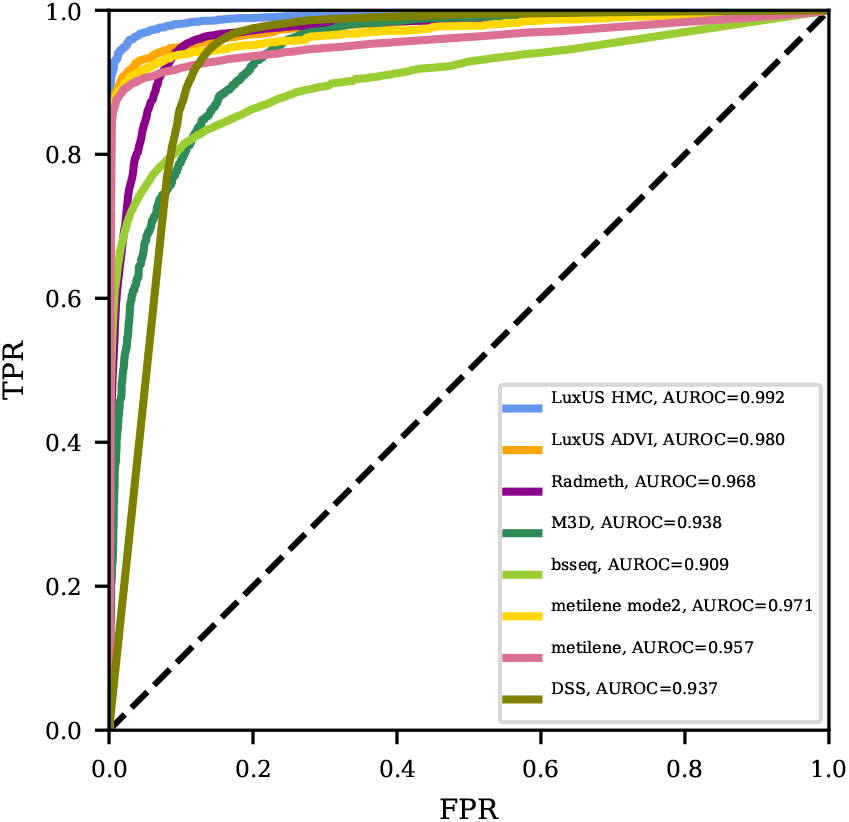
Receiver operating characteristics curves for LuxUS with HMC and ADVI model estimation, RADMeth, M^3^D, bsseq, metilene and DSS for the simulated data by Hebestreit *et al.* (2013). The dashed black line shows the expected ROC curve for random guessing.

## 4 Discussion

The LuxUS method was tested on both simulated and real data sets. The results for the simulated data show that including spatial correlation to the model gives clear advantage when compared to the analysis with only one cytosine being analysed at a time. Comparison with other methods showed that our method performed as well or better than the other methods in almost all simulation settings. In real bisulfite sequencing data analysis LuxUS was more conservative than RADMeth, as the number of differentially methylated cytosines found by LuxUS was considerably lower than for RADMeth. This of course depends on the chosen significance levels and Bayes factor cutoff-values. There was a clear overlap between the differentially methylated cytosines retrieved from these two methods, as three-fourths of the differentially methylated cytosines found by LuxUS were also found by RADMeth. The follow-up analysis with GREAT tool implied that the both methods were able to find biologically significant DMRs.

Based on the results on the simulated data sets, the optimal number of cytosines in a window might be 5 − 10, as this enhances the statistical power of the statistical testing, while the model estimation can be done computationally efficiently. Based on the AUROC values calculated for the simulated data with 50% DMR proportion, both the HMC and ADVI methods for estimating the model seem to perform equally well. However, with 5% DMR proportion the average precision values LuxUS HMC were higher than for LuxUS ADVI. The HMC approach provides better model inference tools for cases where closer investigation on the fitted model is desirable, whereas ADVI is considerably faster. Simple two-group comparison simulation experiments were conducted both with a 50% and a more realistic 5% DMR proportions, which both yielded similar results. Depending on the simulation setting, usually either LuxUS HMC or metilene mode 2 showed best performance. For the simulated data set by Hebestreit *et al.* (2013), all of the methods performed nearly equally well, albeit the preprocessing of the data was done with respect to the restrictions of the RADMeth tool.

Our method enables including both continuous and binary covariates into the fixed effect design matrix and inference on the estimated model, which is not possible with some other tools such as RADMeth, which cannot handle continuous covariates and does not provide a summary of the fitted models. Some tools, such as metilene, cannot take into account any covariates and allow only a comparison between two groups. Accounting for confounding factors is important as several factors, such as age of an individual and smoking history, are known to affect DNA methylation. We demostrated the advantage of this feature with simulation experiments where two confounding covariates were included. In these experiments LuxUS had the best AUROC and average precision values. The metilene (mode 2) tool, which performed well with the simple two-group comparison simulation setting, could not reach the same AUROC and AP levels as the tools which could take the confounding effects into account. With LuxUS, the posterior samples for the linear model coefficients and variance parameters can be explored using the summary and plotting utilities provided in Stan.

Two questions concerning our model are that does the correlation structure match reality well enough and how the priors for the variance parameters should be set. We have provided a set of default parameters for the tool, but the parameter values can be adjusted by the user to match the problem at hand and to the available computational resources. Similarly, the choice of the parameters used in the preanalysis step and the cutoff-value for the Bayes factors affect the final results. As we use the Savage-Dickey Bayes factor estimate, it may be advisable to empirically calibrate Bayes factor cutoffs for significance e.g. by using some known differentially methylated loci. As the ADVI estimates of the posteriors seem to underestimate variance, the Savage-Dickey estimates of the Bayes factors tend to have very high values. To scale down the Bayes factor magnitude, one could opt for using wider bandwidth for the kernel density estimation used in the Savage-Dickey estimation calculation. For example, the Scott’s “rule of thumb” bandwidth which is currently used in the kernel density estimation step could be doubled to produce more conservative kernel density estimates.

Finally, it is possible to extend our model to incorporate the oxidized methylcytosine species into the analysis as done previously in LuxGLM.

## 5 Conclusion

We have proposed a new tool for detecting differential methylation, which uses the spatial correlation of neighboring cytosines to enhance the accuracy of detecting differential methylation. The presented results show that our tool is able to quantify differential methylation from both simulated and real BS-seq data. Comparisons on simulated data show that our model performs as well as, or even better, than previous methods.

The provided preanalysis step can be used to reduce the number of genomic windows for which the Bayesian analysis is done, and the computations can be parallelised for computational efficiency. Opting for variational inference in the model estimation step reduces the needed computation time even further, without having to compromise the accuracy of the fitted model. The tool is available at https://github.com/hallav/LuxUS.

## Acknowledgements

We acknowledge the computational resources provided by the Aalto Science-IT project.

## References

Carpenter, B., Gelman, A., Hoffman,M. D., Lee, D., Goodrich, B., Betancourt, M., Brubaker, M., Guo, J., Li, P., Riddell, A. (2017) Stan: A probabilistic programming language, Journal of Statistical Software, 76(1).

Dolzhenko, E., Smith, A. D. (2014) Using beta-binomial regression for high-precision differential methylation analysis in multifactor whole-genome bisulfite sequencing experiments, BMC Bioinformatics, 15:215.

Eckhardt, F., Lewin, J., Cortese, R., Rakyan, V. K., Attwood, J., Burger, M., Burton, J., Cox, T. V., Davies, R., Down, T. A., Haefliger, C., Horton, R., Howe, K., Jackson, D. K., Kunde, J., Koenig, C., Liddle, J., Niblett, D., Otto, T., Pettett, R., Seemann, S., Thompson, C., West, T., Rogers, J., Olek, A., Berlin, K., Beck, S. (2006) DNA methylation profiling of human chromosomes 6, 20 and 22. Nature Genetics, 38(12), 1378–85.

Feng, H., Conneely, K. N., and Wu, H. (2014). A Bayesian hierarchical model to detect differentially methylated loci from single nucleotide resolution sequencing data. Nucleic acids research, 42(8), e69–e69.

Hansen, K. D., Timp, W., Bravo, H. C., Sabunciyan, S., Langmead, B., McDonald, O. G., Wen, B., Wu, H., Liu, Y., Diep, D., Briem, E., Zhang, K., Irizarry, R. A. and Feinberg, A. P. (2011) Increased methylation variation in epigenetic domains across cancer types. Nature Genetics, 43(8), 768–775.

Hansen, K. D., Langmead, B., Irizarry, R. A. (2012) BSmooth: from whole genome bisulfite sequencing reads to differentially methylated regions. Genome Biology, 13:R83.

Hansen, K. D. (2018). bsseqData: Example whole genome bisulfite data for the bsseq package. R package version 0.20.0.

Hascher, A., Haase, A. K., Hebestreit, K., Rohde, C., Klein, H. U., Rius, M., Jungen, D., Witten, A., Stoll, M., Schulze, I., Ogawa, S., Wiewrodt, R., Tickenbrock, L., Berdel, W. E., Dugas, M., Thoennissen, N. H. and Müller-Tidow, C. (2014) DNA Methyltransferase Inhibition Reverses Epigenetically Embedded Phenotypes in Lung Cancer Preferentially Affecting Polycomb Target Genes. Clinical Cancer Research, 20(4), 814–826.

Hebestreit K., Dugas M., Klein H. (2013) Detection of significantly differentially methylated regions in targeted bisulfite sequencing data. Bioinformatics, 29(13), 1647–1653.

Jühling, F., Kretzmer, H., Bernhart, S. H., Otto, C., Stadler, P. F., and Hoffmann, S. (2016) metilene: Fast and sensitive calling of differentially methylated regions from bisulfite sequencing data. Genome research, 26(2), 256–262.

Kass, R. and Raftery, A. (1995) Bayes factors. Journal of the American Statistical Association. 90 (430): 791.

Korthauer, K., Chakraborty, S., Benjamini, Y., Irizarry, R. A. (2018) Detection and accurate false discovery rate control of differentially methylated regions from whole genome bisulfite sequencing. Biostatistics, kxy007.

Kucukelbir, A., Ranganath, R., Gelman, A., Blei, D. (2015) Automatic Variational Inference in Stan. Advances in Neural Information Processing Systems 28.

Malonzo, M., Halla-aho, V., Konki, M., Lund, R., Lähdesmäki, H. (2018) LuxRep: a technical replicate-aware method for bisulfite sequencing data analysis. bioRxiv 444711 [preprint]. Available from: https://doi.org/10.1101/444711

Mayo, T. R., Schweikert, G., Sanguinetti, G. (2014) M3D: a kernel-based test for spatially correlated changes in methylation profiles. Bioinformatics, 31(6), 809–16.

McLean, C. Y., Bristor, D., Hiller, M., Clarke, S. L., Schaar, B. T., Lowe, C. B., Wenger, A. M., and Bejerano, G. (2010) GREAT improves functional interpretation of cis-regulatory regions. Nature Biotechnology, 28(5), 495–501.

Park, Y., and Wu, H. (2016). Differential methylation analysis for BS-seq data under general experimental design. Bioinformatics, 32(10), 1446–1453.

Rackham, O. J., Langley, S. R., Oates, T., Vradi, E., Harmston, N., Srivastava, P. K., Behmoaras, J., Dellaportas, P., Bottolo, L., Petretto, E. (2017) A Bayesian approach for analysis of whole-Genome bisulfite sequencing data identifies disease-associated changes in DNA methylation. Genetics, 205(4), 1443–1458.

Song, Y., Ren, H., Lei, J. (2017) Collaborations between CpG sites in DNA methylation. International Journal of Modern Physics B, Vol. 31, No. 20, 1750243.

Stan Development Team (2018) CmdStan: the command-line interface to Stan, Version 2.18.0. http://mc-stan.org

Stan Development Team (2017) PyStan: the Python interface to Stan, Version 2.16.0.0. http://mc-stan.org

Virtanen, P., Gommers, R., Oliphant, T.E. et al. (2020) SciPy 1.0: fundamental algorithms for scientific computing in Python. Nature Methods, Vol. 17, 261–272.

Wen, Y., Chen, F., Zhang, Q., Zhuang, Y., Li, Z. (2016) Detection of differentially methylated regions in whole genome bisulfite sequencing data using local Getis-Ord statistics. Bioinformatics, 32(22), 3396–3404.

Äijo, T., Yue, X., Rao, A., Lähdesmäki, H. (2016) LuxGLM: a probabilistic covariate model for quantification of DNA methylation modifications with complex experimental designs. Bioinformatics, 32(17), i511–i519.

