## Supplementary file for "LuxUS: DNA Methylation Analysis Using Generalized Linear Mixed Model with Spatial Correlation"

#### Supplementary information

Viivi Halla-aho and Harri Lähdesmäki

May 4, 2020

#### S1 Methods

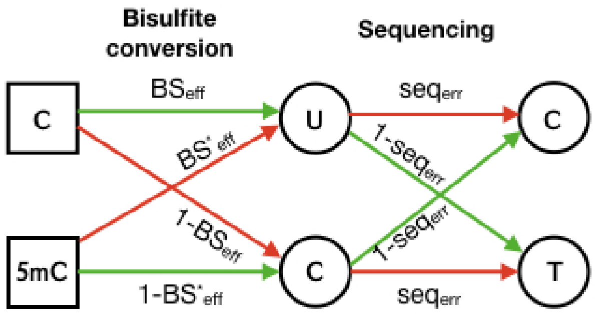


Figure S1: The probability tree for observing a C or T in sequencing data when the true methylation state is methylated or unmethylated.

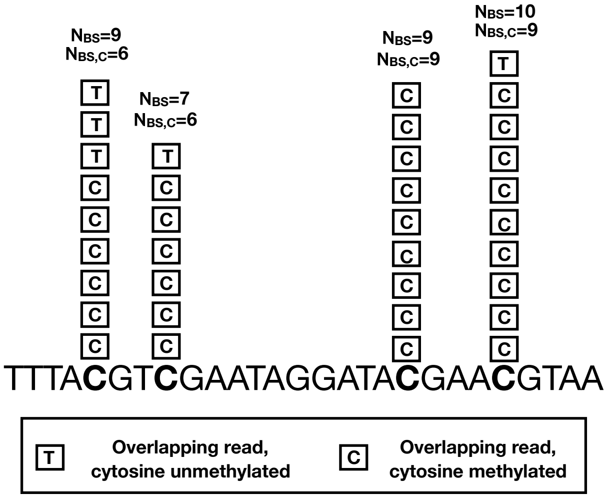


Figure S2: A demonstration of a genomic region with four cytosines with CpG context and the read count data. The squares represent the overlapping reads, from which we get the count data $N_{BS}$ and $N_{BS,C}$, which are the total read count and the count of reads in which the cytosine in question was observed as a C, respectively.

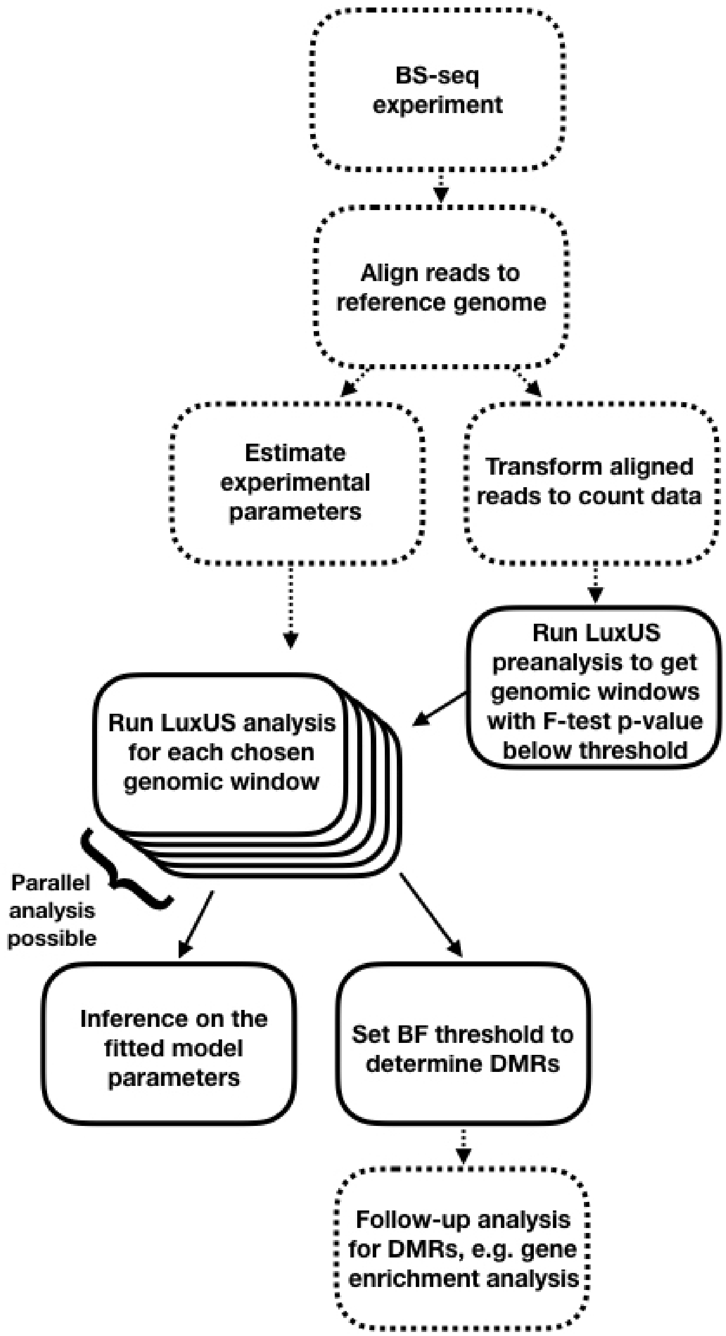


Figure S3: Simplified workflow in LuxUS analysis starting from the bisulfite sequencing and ending in follow-up analysis for the found DMRs. The dashed lines indicate workflow steps that are beyond the scope of LuxUS and which must be performed with another tools.

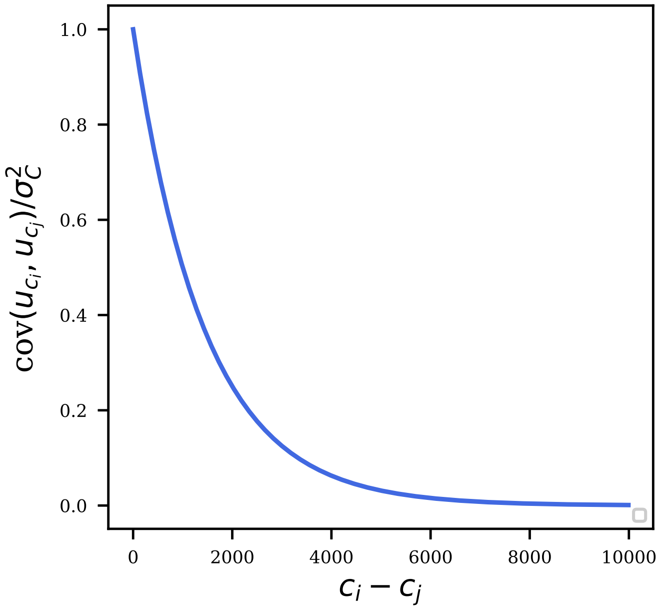


Figure S4: The covariance term from Eq. 10 divided by variance term $\sigma_C^2$, plotted as a function of the genomic distance between two cytosines. The length-scale parameter $\ell$ is set to its prior mean value 38 for the plot. The plot shows how the covariance term decreases as the distance $c_i - c_j$ grows.

#### S1.1 Model estimation

In this section the model estimation methods used in LuxUS are briefly explained. We start from the variational inference (VI) approach and base our description to work by Kucukelbir et al. (2015) and Blei et al. (2017). First, we define the problem of approximating the posterior distribution $p(\theta|\mathbf{X})$, where $\theta$ is a set of latent random variables and $\mathbf{X}$ is a set of observations. We attempt to approximate $p(\theta|\mathbf{X})$ with a simpler distribution $q(\theta;\phi)$, where $\phi$ are the parameters of the approximation.

To find the best approximation $q(\theta;\phi)$ with parameters $\phi$, Kullback-Leibler divergence between the approximation and the true posterior distribution is minimized

$$\min_{\phi} KL(q(\theta;\phi)\|p(\theta|\mathbf{X})). \tag{1}$$

As the above-mentioned KL divergence is often intractable, the evidence lower bound (ELBO) $\mathcal{L}(\phi)$ is maximized instead. Basically, ELBO is the negative KL divergence plus the logarithm of the evidence, $\log p(\mathbf{X})$. The formula for ELBO is

$$\mathcal{L}(\phi) = E_{q(\theta)}[\log p(\mathbf{X},\theta)] - E_{q(\theta)}[\log q(\theta;\phi)]. \tag{2}$$

Maximizing ELBO minimizes the KL divergence. The family of distributions for $q(\theta;\phi)$ is first set and then ELBO is maximized with respect to parameters $\phi$. The result of the optimization is the variational approximation to the posterior distribution $p(\theta|\mathbf{X})$. Choosing the approximative family often requires careful studying of the distribution to be approximated.

Probabilistic programming language Stan, which we use to implement LuxUS, uses automatic differentation variational inference (ADVI) algorithm for variational inference. The basic idea in ADVI is to first transform the latent variables into real coordinate space and then choose Gaussian variational distribution family to approximate the posterior. Choosing Gaussian approximation in the transformed space means that the approximation is non-Gaussian after it has been transformed back to the original space. This trick enables automation of variational inference. The objective of minimization Kullback-Leibler (KL) distance between the true and approximate posterior distributions is then achieved by combining automatic differentiation and stochastic optimization. In LuxUS we have utilized the mean-field algorithm feature of ADVI. In mean-field version of the ADVI algorithm, the variational approximation to the transformed parameters is Gaussian and fully factorised. This means that the latent variables are independent of each other and each of them has a variational factor of their own in $q(\theta;\phi)$.

While in the variational inference approach the problem of approximating posterior distribution $p(\theta|\mathbf{X})$ was tackled by solving an optimization problem, in the Markov chain Monte Carlo (MCMC) approaches we sample from a Markov chain which is bound to converge to the posterior distribution. Here we briefly describe the Hamiltonian Monte Carlo sampling method, which is used in Stan. We use Gelman et al. (2014) as reference for this section. One of the simple

MCMC sampling methods is Metropolis algorithm, in which the samples are retrieved form a random walk with an acceptance rule that guarantees asymptotic convergence to the posterior distribution. Hamiltonian Monte Carlo is a sophisticated version of the Metropolis algorithm, where the random walk behavior is made more efficient by introducing physics-inspired momentum variables $\phi$. The momentum variables are updated together with the sampled parameters $\theta$ during the sampling.

By default, Stan uses no-U-turn-sampler (NUTS), which is a locally adaptive version of HMC. In NUTS the number of steps per iteration is adapted for every iteration, allowing the sampler to explore the posterior space efficiently until the trajectory turns around. This prevents the sampler from circling around in the posterior space.

MCMC methods are often computationally more intensive than VI methods, which can take advantage of efficient optimization algorithms. As a downside, VI methods can only find an approximation which is close to the target distribution but convergence to the true posterior distribution is not guaranteed, whereas MCMC methods guarantee asymptotic convergence to the target distribution. (Blei et al., 2017)

### S2 Analysis of colon cancer WGBS-seq data

#### S2.1 Preanalysis and setting priors

The LuxUS model was tested using a WGBS-seq data set, consisting of matched human colon and colon cancer samples. The processed sequencing data was provided for two chromosomes, 21 and 22. First, the genomic windows for LuxUS analysis were determined using the preanalysis method. The p-value cutoff for the preanalysis F-test for the significance of the cancer covariate was set to 0.1. The p-value cutoff value was chosen so that it would not be too conservative and filter out too many potentially differentially methylated cytosines. For a cytosine to be added in a genomic window it had to have at least coverage of one in at least one sample from both case and control groups. Additionally, the mean coverage over the genomic window had to be at least 5 at least in one sample from both case and control groups for the genomic window to be accepted for analysis. The maximum number of cytosines in a genomic window was set to 20 and the maximum length of the genomic window was set to 2000 basepairs. This resulted in 9945 genomic windows that passed the preanalysis phase. The hyperparameters for the random effect and noice variances were set to $\alpha_R = 2$, $\beta_R = 2$, $\alpha_C = 2$, $\beta_C = 2$, $\alpha_E = 2$ and $\beta_E = 2$. The variance parameter for the coefficients $\mathbf{b}$, $\sigma_b^2$, was set to 15. The bisulfite conversion efficiency, the incorrect bisulfite conversion efficiency and sequencing error were set to $\text{BS}_{\text{eff}} = 1$, $\text{BS}^*_{\text{eff}} = 0$ and $\text{seq}_{\text{err}} = 0$ respectively for each sample, which correspond to perfect bisulfite conversion efficiency, perfect incorrect bisulfite conversion efficiency and to no sequencing error. This way the results are comparable to RADMeth tool, which cannot take the experimental parameters into account.

#### S2.2 Results

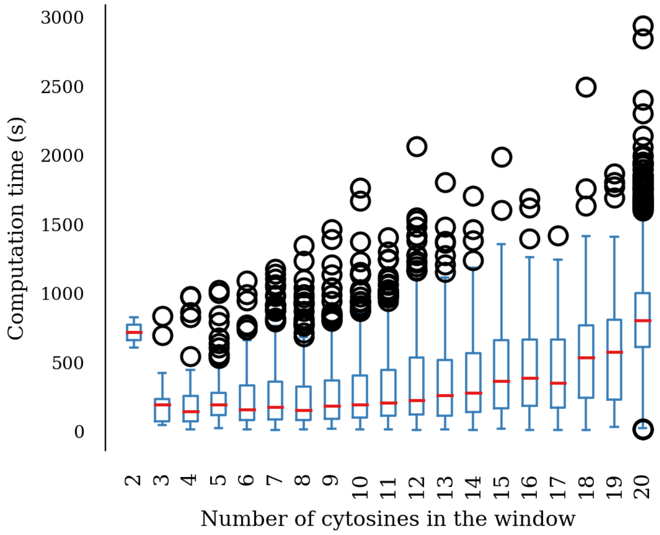


Figure S5: Boxplots of the computation times for the genomic windows for the colon cancer WGBS data. One boxplot represents all the windows with the same number of cytosines in the genomic window.

| Ontology | # Term Name | Hyper Rank | Hyper Raw P-Value | Hyper FDR Q-Val | Hyper Fold Enrichment | Hyper Foreground Region Hits | Hyper Total Regions | Hyper Region Set Coverage | Hyper Foreground Gene Hits | Total Genes Annotated |
|---|---|---|---|---|---|---|---|---|---|---|
| Disease Ontology | partial epilepsy | 1 | 3.93334e-178 | 8.79101e-175 | 3.6243 | 681 | 8525 | 3.22% | 2 | 58 |
| | temporal lobe epilepsy | 1 | 3.93334e-178 | 8.79101e-175 | 3.6243 | 681 | 8525 | 3.22% | 2 | 48 |
| | oligodendroglioma | 3 | 1.02499e-151 | 7.63616e-149 | 3.1820 | 692 | 9867 | 3.28% | 2 | 18 |
| | epilepsy | 5 | 6.05340e-104 | 2.70587e-101 | 2.4017 | 759 | 14338 | 3.59% | 4 | 198 |
| | T-cell leukemia | 6 | 7.50644e-93 | 2.79615e-90 | 6.2237 | 200 | 1458 | 0.95% | 1 | 60 |
| | leishmaniasis | 6 | 7.50644e-93 | 2.79615e-90 | 6.2237 | 200 | 1458 | 0.95% | 1 | 21 |
| | visceral leishmaniasis | 6 | 7.50644e-93 | 2.79615e-90 | 6.2237 | 200 | 1458 | 0.95% | 1 | 9 |
| | nasopharynx carcinoma | 9 | 1.18992e-90 | 2.95498e-88 | 2.1238 | 859 | 18351 | 4.07% | 3 | 194 |
| | microcytic anemia | 14 | 3.20167e-74 | 5.11124e-72 | 4.8838 | 200 | 1858 | 0.95% | 1 | 6 |
| | iron deficiency anemia | 14 | 3.20167e-74 | 5.11124e-72 | 4.8838 | 200 | 1858 | 0.95% | 1 | 2 |
| | deficiency anemia | 14 | 3.20167e-74 | 5.11124e-72 | 4.8838 | 200 | 1858 | 0.95% | 1 | 2 |
| | nutritional deficiency disease | 14 | 3.20167e-74 | 5.11124e-72 | 4.8838 | 200 | 1858 | 0.95% | 1 | 18 |
| | endometrial adenocarcinoma | 18 | 4.91903e-69 | 6.10779e-67 | 3.0917 | 326 | 4784 | 1.54% | 2 | 31 |
| | ependymoma | 19 | 4.50544e-59 | 5.29982e-57 | 3.5028 | 233 | 3018 | 1.10% | 1 | 12 |
| | chronic granulomatous disease | 23 | 1.07203e-47 | 1.04173e-45 | 5.1606 | 120 | 1055 | 0.57% | 1 | 14 |
| | spondyloarthropathy | 24 | 2.88504e-47 | 2.68669e-45 | 2.0350 | 492 | 10969 | 2.33% | 1 | 73 |
| | spondylitis | 24 | 2.88504e-47 | 2.68669e-45 | 2.0350 | 492 | 10969 | 2.33% | 1 | 64 |
| | ankylosing spondylitis | 24 | 2.88504e-47 | 2.68669e-45 | 2.0350 | 492 | 10969 | 2.33% | 1 | 64 |
| | Euglenozoa infectious disease | 29 | 1.74247e-41 | 1.34291e-39 | 3.0177 | 200 | 3007 | 0.95% | 1 | 39 |
| | placental abruption | 30 | 3.52838e-38 | 2.62864e-36 | 2.3719 | 280 | 5356 | 1.33% | 1 | 14 |
| | kidney neoplasm | 32 | 1.08598e-31 | 7.58489e-30 | 2.1653 | 280 | 5867 | 1.33% | 1 | 56 |
| | renal malignant neoplasm | 32 | 1.08598e-31 | 7.58489e-30 | 2.1653 | 280 | 5867 | 1.33% | 1 | 40 |
| | cleft lip | 34 | 1.95918e-31 | 1.28787e-29 | 2.1572 | 280 | 5889 | 1.33% | 1 | 54 |
| | pilocytic astrocytoma | 37 | 4.38232e-28 | 2.64716e-26 | 2.2199 | 233 | 4762 | 1.10% | 1 | 12 |
| | malignant melanoma of conjunctiva | 39 | 2.57495e-22 | 1.47565e-20 | 3.2457 | 93 | 1300 | 0.44% | 1 | 3 |
| | nephroblastoma | 39 | 2.57495e-22 | 1.47565e-20 | 3.2457 | 93 | 1300 | 0.44% | 1 | 70 |
| | hereditary Wilms' cancer | 39 | 2.57495e-22 | 1.47565e-20 | 3.2457 | 93 | 1300 | 0.44% | 1 | 54 |
| | leukocyte adhesion deficiency | 42 | 8.11164e-19 | 4.31655e-17 | 2.7497 | 100 | 1650 | 0.47% | 1 | 5 |
| | intermediate coronary syndrome | 42 | 8.11164e-19 | 4.31655e-17 | 2.7497 | 100 | 1650 | 0.47% | 1 | 22 |
| | oxyphilic adenoma | 83 | 3.39523e-6 | 9.14258e-5 | 2.2896 | 38 | 753 | 0.18% | 1 | 15 |
| | leukoencephalopathy | 86 | 9.86362e-6 | 2.56340e-4 | 2.1351 | 40 | 850 | 0.19% | 1 | 21 |

| | |
|---|---|
| The test set contains 21,127 (2%) of all 958,541 regions. | The test set picked 282 genes, the background set picked 600 genes. |
| *Disease Ontology* has 2,235 terms covering 7,886 (44%) of all 18,041 genes. | 2,235 ontology terms were tested (100%) using an annotation count range of [1, Inf]. |

| |
|---|
| GREAT version 3.0.0 |
| Species assembly: hg19 |
| Association rule: Basal+extension: 5000 bp upstream, 1000 bp downstream, 1000000 bp max extension, curated regulatory domains included |

Figure S6: GREAT results table for the Disease Ontology for results from LuxUS analysis.

| Ontology | # Term Name | Hyper Rank | Hyper Raw P-Value | Hyper FDR Q-Val | Hyper Fold Enrichment | Hyper Foreground Region Hits | Hyper Total Regions | Hyper Region Set Coverage | Hyper Foreground Gene Hits | Total Genes Annotated |
|---|---|---|---|---|---|---|---|---|---|---|
| MSigDB Perturbation | Genes down-regulated in Cockayne syndrome fibroblasts rescued by expression of ERCC6 [GeneID=2074] off a plasmid vector. | 1 | 0.00000 | 0.00000 | 2.9147 | 2489 | 38744 | 11.78% | 2 | 36 |
| | Genes up-regulated in metastatic vs non-metastatic HNSCC (head and neck squamous cell carcinoma) samples. | 1 | 0.00000 | 0.00000 | 2.6299 | 2195 | 37868 | 10.39% | 2 | 325 |
| | Genes in the most frequently homozygous deleted loci in a panel of glioma cell lines. | 1 | 0.00000 | 0.00000 | 2.7615 | 2175 | 35735 | 10.29% | 1 | 50 |
| | Up-regulated genes distinguishing between two subtypes of gastric cancer: advanced (AGC) and early (EGC). | 1 | 0.00000 | 0.00000 | 2.6323 | 3154 | 54363 | 14.93% | 4 | 170 |
| | Genes down-regulated in mucinous ovarian carcinoma tumors of grades 1 and 2 compared to the normal ovarian survace epithelium tissue. | 1 | 0.00000 | 0.00000 | 4.1435 | 1453 | 15910 | 6.88% | 2 | 63 |
| | Genes down-regulated in CD-1 compared to CD-2 cluster of multiple myeloma samples. | 1 | 0.00000 | 0.00000 | 2.6263 | 2192 | 37867 | 10.38% | 2 | 51 |
| | Genes down-regulated in follicular epithelial stem cells after transgenic expression of GR [GeneID=2908] under control of the keratin5 (K5) [GeneID=3852] promoter. | 7 | 7.92241e-317 | 3.80729e-314 | 2.4340 | 2195 | 40916 | 10.39% | 2 | 120 |
| | Cluster P4 of genes with similar expression profiles in peripheral T lymphocytes after FOXP3 [GeneID=50943] loss of function (LOF). | 8 | 8.71517e-297 | 3.66473e-294 | 4.0005 | 1002 | 11364 | 4.74% | 2 | 97 |
| | Genes down-regulated in a breast cancer cell line resistant to tamoxifen [PubChem=5376] compared to the parental line sensitive to the drug. | 9 | 5.66914e-266 | 2.11900e-263 | 4.4293 | 802 | 8215 | 3.80% | 1 | 52 |
| | Cluster A: genes up-regulated in primary lung tumors driven by KRAS [GeneID=3845] activation and loss of STK11 [GeneID=6794]; also up-regulated in human squamous cell carcinoma (SCC) vs adenocarcinoma subtype of NSCLC (non-small cell lung cancer). | 9 | 5.66914e-266 | 2.11900e-263 | 4.4293 | 802 | 8215 | 3.80% | 1 | 12 |
| | Genes down-regulated by trabectedin [PubChem=3199] and its synthetic analog phthalascidin Pt 650 in HCT116 cells (colon cancer). | 9 | 5.66914e-266 | 2.11900e-263 | 4.4293 | 802 | 8215 | 3.80% | 1 | 50 |
| | Genes corresponding to the histamine [PubChem=774] response network. | 9 | 5.66914e-266 | 2.11900e-263 | 4.4293 | 802 | 8215 | 3.80% | 1 | 32 |
| | Cluster 5: genes changed in primary keratinocytes by UVB irradiation. | 9 | 5.66914e-266 | 2.11900e-263 | 4.4293 | 802 | 8215 | 3.80% | 1 | 46 |
| | Genes up-regulated by estradiol [PubChem=5757] and not modulated by ESRRA [GeneID=2101] in MCF-7 cells (breast cancer). | 9 | 5.66914e-266 | 2.11900e-263 | 4.4293 | 802 | 8215 | 3.80% | 1 | 18 |
| | Genes up-regulated in U2OS cells (osteosarcoma) upon knockdown of both HDAC1 and HDAC2 [GeneID=3065;3066] by RNAi. | 15 | 3.63284e-265 | 8.14725e-263 | 2.1886 | 2282 | 47307 | 10.80% | 3 | 224 |
| | Cluster 5 of aberrantly hypomethylated genes in blasts from AML (acute myeloid leukemia) patients. | 16 | 9.52687e-260 | 2.00302e-257 | 3.7467 | 950 | 11504 | 4.50% | 3 | 48 |
| | Genes discriminating between direct (cisplatin, MMS, mitomycin C [PubChem=2767;4156;5746]) and indirect (paclitaxel, hydroxyurea, etoposide [PubChem=4666;3657;36462]) acting genotoxins at 4 h time point. | 17 | 1.50051e-255 | 2.96924e-253 | 3.3614 | 1075 | 14510 | 5.09% | 2 | 36 |
| | Genes up-regulated in B lymphocytes from patients with primary immunodefiency syndrom. | 19 | 2.84995e-238 | 5.04592e-236 | 3.8568 | 842 | 9905 | 3.99% | 2 | 47 |
| | Genes up-regulated by expression of GAB2 [GeneID=9846] in K562 cells (chronic myeloid leukemia (CML) cell line with p210 BCR-ABL [GeneID=613;25]). | 20 | 2.80123e-237 | 4.71167e-235 | 4.7158 | 670 | 6446 | 3.17% | 1 | 31 |
| | Genes up-regulated in primary keratinocytes by expression of p50 (NFKB1) and p65 (RELA) [GeneID=4790;5970] components of NFKB. | 22 | 2.74869e-228 | 4.20300e-226 | 3.3977 | 948 | 12659 | 4.49% | 4 | 91 |

| | |
|---|---|
| The test set contains 21,127 (2%) of all 958,541 regions. | The test set picked 282 genes, the background set picked 600 genes. |
| *MSigDB Perturbation* has 3,364 terms covering 17,091 (95%) of all 18,041 genes. | 3,364 ontology terms were tested (100%) using an annotation count range of [1, Inf]. |
| GREAT version 3.0.0 | |
| Species assembly: hg19 | |
| Association rule: Basal+extension: 5000 bp upstream, 1000 bp downstream, 1000000 bp max extension, curated regulatory domains included | |

Figure S7: GREAT results table for the MSigDB Perturbations ontology for results from LuxUS analysis.

| Ontology | # Term Name | Hyper Rank | Hyper Raw P-Value | Hyper FDR Q-Val | Hyper Fold Enrichment | Hyper Foreground Region Hits | Hyper Total Regions | Hyper Region Set Coverage | Hyper Foreground Gene Hits | Total Genes Annotated |
|---|---|---|---|---|---|---|---|---|---|---|
| Disease Ontology | oligodendroglioma | 1 | 0.00000 | 0.00000 | 2.4353 | 3438 | 9867 | 2.51% | 3 | 18 |
| | ependymoma | 6 | 7.69128e-226 | 2.86500e-223 | 2.6517 | 1145 | 3018 | 0.83% | 1 | 12 |
| | contagious pustular dermatitis | 10 | 8.06614e-171 | 1.80278e-168 | 2.0776 | 1469 | 4942 | 1.07% | 1 | 26 |
| | microcytic anemia | 15 | 2.78267e-116 | 4.14618e-114 | 2.4903 | 662 | 1858 | 0.48% | 1 | 6 |
| | iron deficiency anemia | 15 | 2.78267e-116 | 4.14618e-114 | 2.4903 | 662 | 1858 | 0.48% | 1 | 2 |
| | deficiency anemia | 15 | 2.78267e-116 | 4.14618e-114 | 2.4903 | 662 | 1858 | 0.48% | 1 | 2 |
| | nutritional deficiency disease | 15 | 2.78267e-116 | 4.14618e-114 | 2.4903 | 662 | 1858 | 0.48% | 1 | 18 |
| | leukocyte adhesion deficiency | 23 | 2.46485e-97 | 2.39520e-95 | 2.4399 | 576 | 1650 | 0.42% | 1 | 5 |
| | intermediate coronary syndrome | 23 | 2.46485e-97 | 2.39520e-95 | 2.4399 | 576 | 1650 | 0.42% | 1 | 22 |
| | T-cell leukemia | 33 | 2.90461e-78 | 1.96721e-76 | 2.3634 | 493 | 1458 | 0.36% | 1 | 60 |
| | leishmaniasis | 33 | 2.90461e-78 | 1.96721e-76 | 2.3634 | 493 | 1458 | 0.36% | 1 | 21 |
| | visceral leishmaniasis | 33 | 2.90461e-78 | 1.96721e-76 | 2.3634 | 493 | 1458 | 0.36% | 1 | 9 |

| | |
|---|---|
| The test set contains 137,142 (14%) of all 958,541 regions. | The test set picked 595 genes, the background set picked 600 genes. |

Figure S8: GREAT results table for the Disease Ontology for results from RAD-Meth analysis.

| Ontology | # Term Name | Hyper Rank | Hyper Raw P-Value | Hyper FDR Q-Val | Hyper Fold Enrichment | Hyper Foreground Region Hits | Hyper Total Regions | Hyper Region Set Coverage | Hyper Foreground Gene Hits | Total Genes Annotated |
|---|---|---|---|---|---|---|---|---|---|---|
| MSigDB Perturbation | Genes down-regulated in T47D cells (breast cancer) after COBRA1 [GeneID=25920] knockdown by RNAi. | 1 | 0.00000 | 0.00000 | 2.1231 | 3109 | 10235 | 2.27% | 1 | 29 |
| | Genes down-regulated in a breast cancer cell line resistant to tamoxifen [PubChem=5376] compared to the parental line sensitive to the drug. | 1 | 0.00000 | 0.00000 | 2.1611 | 2540 | 8215 | 1.85% | 1 | 52 |
| | Down-regulated genes in hepatocellular carcinoma (HCC) subclass G3, defined by unsupervised clustering. | 1 | 0.00000 | 0.00000 | 2.2523 | 3603 | 11181 | 2.63% | 3 | 49 |
| | Genes down-regulated in primary fibroblast cell culture point after infection with HCMV (AD169 strain) at 6 h time point that were not down-regulated at the previous time point, 4 h. | 1 | 0.00000 | 0.00000 | 2.1366 | 12865 | 42085 | 9.38% | 3 | 157 |
| | Cluster 3: genes maximally expressed at 8 hr time point during differentiation of 3T3-L1 fibroblasts into adipocytes in response to adipogenic hormones. | 1 | 0.00000 | 0.00000 | 2.1577 | 4681 | 15163 | 3.41% | 2 | 39 |
| | Genes down-regulated in follicular epithelial stem cells after transgenic expression of GR [GeneID=2908] under control of the keratin5 (K5) [GeneID=3852] promoter. | 1 | 0.00000 | 0.00000 | 2.4479 | 14330 | 40916 | 10.45% | 3 | 120 |
| | Genes down-regulated in NIH3T3 cells (fibroblasts) transformed by activated KRAS [GeneID=3845] vs those reverted to normal cells upon over-expression of a dominant negative form of CDC25 [GeneID=5923]. | 1 | 0.00000 | 0.00000 | 2.3568 | 3029 | 8983 | 2.21% | 1 | 50 |
| | Selected genes down-regulated during invasion of lymphatic vessels during metastasis. | 1 | 0.00000 | 0.00000 | 2.2583 | 3862 | 11953 | 2.82% | 3 | 36 |
| | Genes up-regulated in ANBL-6 cell line (multiple myeloma, MM) co-cultured with bone marrow stromal cells compared to those grown in the presence of IL6 [GeneID=3569]. | 1 | 0.00000 | 0.00000 | 2.3568 | 3029 | 8983 | 2.21% | 1 | 59 |
| | Genes most strongly up-regulated in kidney glomeruli isolated from TCF21 [GeneID=6943] knockout mice. | 1 | 0.00000 | 0.00000 | 2.1328 | 4132 | 13541 | 3.01% | 3 | 36 |
| | Cluster 5 of aberrantly hypomethylated genes in blasts from AML (acute myeloid leukemia) patients. | 1 | 0.00000 | 0.00000 | 2.0724 | 3411 | 11504 | 2.49% | 3 | 48 |
| | Genes up-regulated in glioblastoma cell lines displaying spherical growth (cluster-1) compared to those displaying semiadherent or adherent growth phenotype (cluster-2). | 1 | 0.00000 | 0.00000 | 2.5512 | 1917 | 5252 | 1.40% | 2 | 21 |
| | Genes down-regulated in 3T3 cells (fibroblast) upon activation of JNK pathway. | 1 | 0.00000 | 0.00000 | 2.3390 | 3633 | 10856 | 2.65% | 2 | 40 |
| | Genes from the brain cancer stem (cancer stem cell, CSC) signature. | 1 | 0.00000 | 0.00000 | 2.1751 | 3650 | 11729 | 2.66% | 3 | 43 |
| | Genes down-regulated in H358 cells (lung cancer) by inducible expression of FOXA2 [GeneID=3170] in a Tet-off system. | 1 | 0.00000 | 0.00000 | 2.1505 | 3055 | 9929 | 2.23% | 2 | 36 |
| | Genes up-regulated during transition from L0 (non-tumor, not infected with HCV) to L1 (non-tumor, infected with HCV) in the development of hepatocellular carcinoma. | 1 | 0.00000 | 0.00000 | 2.3301 | 2823 | 8468 | 2.06% | 1 | 17 |
| | Genes down-regulated in MEF cells (embryonic fibroblast) upon Cre-lox knockout of DNMT1 [GeneID=1786]. | 1 | 0.00000 | 0.00000 | 2.3390 | 3633 | 10856 | 2.65% | 2 | 25 |
| | Genes up-regulated during epithelial to mesenchymal transition (EMT) induced by TGFB1 [GeneID=7040] in the EpH4 cells (mammary epithelium cell line transformed by HRAS [GeneID=3265]). | 1 | 0.00000 | 0.00000 | 2.3390 | 3633 | 10856 | 2.65% | 2 | 70 |
| | Cluster A: genes up-regulated in primary lung tumors driven by KRAS [GeneID=3845] activation and loss of STK11 [GeneID=6794]; also up-regulated in human squamous cell carcinoma (SCC) vs adenocarcinoma subtype of NSCLC (non-small cell lung cancer). | 1 | 0.00000 | 0.00000 | 2.1611 | 2540 | 8215 | 1.85% | 1 | 12 |
| | Genes up-regulated in osteoblasts from wild type male mice compared to those with AR [GeneID=367] knockout. | 1 | 0.00000 | 0.00000 | 2.3568 | 3029 | 8983 | 2.21% | 1 | 17 |
| | Genes up-regulated similarly in primary fibroblast cultures from Werner syndrom patients and normal old donors compared to those from normal young donors. | 1 | 0.00000 | 0.00000 | 2.0710 | 3169 | 10695 | 2.31% | 2 | 93 |
| | Genes down-regulated in serrated vs conventional colorectal carcinoma (CRC) samples. | 1 | 0.00000 | 0.00000 | 2.0968 | 9103 | 30344 | 6.64% | 3 | 74 |
| | Down-regulated genes from the 324 genes identified by two analytical methods as changed in the mammary tumors induced by transgenic expression of ERBB2 [GeneID=2064]. | 1 | 0.00000 | 0.00000 | 2.0046 | 5643 | 19675 | 4.11% | 5 | 146 |
| | Downregulated in the neocortex of aged adult mice (30-month) vs young adult (5-month) | 1 | 0.00000 | 0.00000 | 2.0647 | 3074 | 10406 | 2.24% | 3 | 79 |
| | Top genes associated with unfavorable survival after surgery of patients with epithelioid mesothelioma. | 1 | 0.00000 | 0.00000 | 2.9695 | 1812 | 4265 | 1.32% | 1 | 12 |
| | Cluster 4: genes with similar expression profiles across follicular thyroid carcinoma (FTC) samples. | 1 | 0.00000 | 0.00000 | 2.1231 | 3109 | 10235 | 2.27% | 1 | 12 |
| | NF-kB-controlled genes down-regulated in endothelial cells in response to viral GPCR protein. | 1 | 0.00000 | 0.00000 | 2.6963 | 8433 | 21860 | 6.15% | 1 | 11 |
| | Down-regulated genes in the expression signature of direct and paracrine viral GPCR signaling in endothelial cells. | 1 | 0.00000 | 0.00000 | 2.2910 | 9008 | 27482 | 6.57% | 3 | 49 |
| | Genes down-regulated by trabectedin [PubChem=3199] and its synthetic analog phthalascidin Pt 650 in HCT116 cells (colon cancer). | 1 | 0.00000 | 0.00000 | 2.1611 | 2540 | 8215 | 1.85% | 1 | 50 |
| | Genes with high-CpG-density promoters (HCP) bearing histone H3 K27 trimethylation mark (H327me3) in embryonic stem cells (ES). | 1 | 0.00000 | 0.00000 | 2.5950 | 2329 | 6273 | 1.70% | 1 | 40 |
| | Genes with low-CpG-density promoters (LCP) bearing the tri-methylation mark at H3K4 (H3K4me3) in MCV6 cells (embryonic fibroblasts trapped in a differentiated state). | 1 | 0.00000 | 0.00000 | 2.3568 | 3029 | 8983 | 2.21% | 1 | 153 |
| | Top genes up-regulated in mesenchyme stem cells (MSC) grown in a tumor conditioned medium, which leads to carcinoma-associated fibroblast phenotype. | 1 | 0.00000 | 0.00000 | 2.3390 | 3633 | 10856 | 2.65% | 2 | 25 |
| | Selected genes up-regulated in WT-A10 cells (gastric epithelium) expressing the H. pilori virulence gene CagA. | 1 | 0.00000 | 0.00000 | 2.3390 | 3633 | 10856 | 2.65% | 2 | 24 |
| | Top 100 probe sets contrubuting to the positive side of the 2nd principal component; associated with adipocytic differentiation. | 1 | 0.00000 | 0.00000 | 2.0824 | 3024 | 10150 | 2.21% | 2 | 87 |
| | Genes down-regulated in Cockayne syndrome fibroblasts rescued by expression of ERCC6 [GeneID=2074] off a plasmid vector. | 1 | 0.00000 | 0.00000 | 2.7338 | 15154 | 38744 | 11.05% | 2 | 36 |
| | Genes down-regulated in neutrophils upon treatment with activated protein C (PROC) [GeneID=5624] of pulmonary inflammation induced by bacterial lipopolysaccharide (LPS). | 1 | 0.00000 | 0.00000 | 2.5020 | 8516 | 23790 | 6.21% | 2 | 27 |
| | Up-regulated genes in angioimmunoblastic lymphoma (AILT) compared to normal T lymphocytes. | 1 | 0.00000 | 0.00000 | 2.3145 | 3971 | 11992 | 2.90% | 2 | 201 |
| | Genes corresponding to the histamine [PubChem=774] response network. | 1 | 0.00000 | 0.00000 | 2.1611 | 2540 | 8215 | 1.85% | 1 | 32 |
| | Genes up-regulated in metastatic vs non-metastatic HNSCC (head and neck squamous cell carcinoma) samples. | 1 | 0.00000 | 0.00000 | 2.6674 | 14452 | 37868 | 10.54% | 4 | 325 |
| | Genes in the most frequently homozygous deleted loci in a panel of glioma cell lines. | 1 | 0.00000 | 0.00000 | 2.7797 | 14212 | 35735 | 10.36% | 1 | 50 |
| | Genes down-regulated in fibroblasts from MLL [GeneID=4297] knockout mice. | 1 | 0.00000 | 0.00000 | 2.3390 | 3633 | 10856 | 2.65% | 2 | 33 |
| | Genes up-regulated in U2OS cells (osteosarcoma) upon knockdown of both HDAC1 and HDAC2 [GeneID=3065;3066] by RNAi. | 1 | 0.00000 | 0.00000 | 2.2416 | 15172 | 47307 | 11.06% | 5 | 224 |
| | Genes up-regulated in pancreatic islets upon knockout of HNF1A [GeneID=6927]. | 1 | 0.00000 | 0.00000 | 2.0653 | 6181 | 20918 | 4.51% | 5 | 163 |
| | Cluster 5: genes changed in primary keratinocytes by UVB irradiation. | 1 | 0.00000 | 0.00000 | 2.1611 | 2540 | 8215 | 1.85% | 1 | 46 |
| | Genes up-regulated by estradiol [PubChem=5757] and not modulated by ESRRA [GeneID=2101] in MCF-7 cells (breast cancer). | 1 | 0.00000 | 0.00000 | 2.1611 | 2540 | 8215 | 1.85% | 1 | 18 |
| | Genes up-regulated in response to both single dose and fractionated radiation that were common to all three cell lines studied. | 1 | 0.00000 | 0.00000 | 2.3390 | 3633 | 10856 | 2.65% | 2 | 32 |
| | Genes showing decreasing expression in brown preadipocytes with increasing ability of the cells to differentiate. | 1 | 0.00000 | 0.00000 | 2.0621 | 3035 | 10287 | 2.21% | 2 | 46 |
| | Genes down-regulated in 9.5 days post coitus (dpc) embryos with COMMD1 [GeneID=150684] knockout compared to normal 9.5 dpc embryos. | 1 | 0.00000 | 0.00000 | 2.1231 | 3109 | 10235 | 2.27% | 1 | 39 |
| | Up-regulated genes distinguishing between two subtypes of gastric cancer: advanced (AGC) and early (EGC). | 1 | 0.00000 | 0.00000 | 2.3648 | 18393 | 54363 | 13.41% | 5 | 170 |
| | Genes up-regulated in MMEC cells (myometrial endothelium) at 6 h after VEGFA [GeneID=7422] stimulation. | 1 | 0.00000 | 0.00000 | 2.0937 | 3420 | 11417 | 2.49% | 3 | 59 |
| | Genes up-regulated in a linear fashion in CD34+ [GeneID=947] cells upon increasing activity levels of STAT5A [GeneID=6776]; predominant long-term growth and self-renewal phenotype. | 1 | 0.00000 | 0.00000 | 2.3568 | 3029 | 8983 | 2.21% | 1 | 59 |
| | Genes up-regulated in MFCF-7 cells (breast cancer) by exogenous HG1 [GeneID=2688]. | 1 | 0.00000 | 0.00000 | 2.4756 | 2728 | 7702 | 1.99% | 2 | 79 |
| | Genes down-regulated in the uteri of ovariectomized mice 6 h after progesterone [PubChem=5994] injection: HOXA10 [GeneID=3206] knockout vs wild type animals. | 1 | 0.00000 | 0.00000 | 2.1231 | 3109 | 10235 | 2.27% | 1 | 18 |
| | Genes co-regulated in uterus during a time course response to progesterone [PubChem=5994]: SOM cluster 12. | 1 | 0.00000 | 0.00000 | 2.0498 | 3744 | 12766 | 2.73% | 2 | 81 |

Figure S9: Part 1 out of 2 of GREAT results table for the MSigDB Perturbations ontology for results from RADMeth analysis.

| | | | | | | | | | | |
|---|---|---|---|---|---|---|---|---|---|---|
| | Genes co-regulated in uterus during a time course response to progesterone [PubChem=5994]: SOM cluster 16. | 1 | 0.00000 | 0.00000 | 2.0222 | 6129 | 21184 | 4.47% | 5 | 78 |
| | Genes commonly up-regulated in CD-1 and CD-2 clusters of multiple myeloma samples and which were higher expressed in the CD-1 group. | 1 | 0.00000 | 0.00000 | 2.2546 | 2307 | 7152 | 1.68% | 2 | 79 |
| | Genes down-regulated in CD-1 compared to CD-2 cluster of multiple myeloma samples. | 1 | 0.00000 | 0.00000 | 2.6625 | 14425 | 37867 | 10.52% | 2 | 51 |
| | Genes down-regulated in glioblastoma cell lines displaying spherical growth (cluster-1) compared to those displaying semiadherent or adherent growth phenotype (cluster-2). | 101 | 2.04502e-316 | 6.81134e-315 | 2.1987 | 2375 | 7550 | 1.73% | 1 | 25 |
| | Genes changed by expression of activated form of HRas[GeneID=3265] in MEF cells (embryonic fibroblast) with or without p65/c-Rel complex [GeneID=5970;5966]. | 101 | 2.04502e-316 | 6.81134e-315 | 2.1987 | 2375 | 7550 | 1.73% | 1 | 22 |
| | Genes up-regulated in malignant skin tumors (squamous cell carcinoma, SCC) formed by treatment with DMBA and TPA [PubChem=6001;4792] in the two stage skin carcinogenesis model. | 101 | 2.04502e-316 | 6.81134e-315 | 2.1987 | 2375 | 7550 | 1.73% | 1 | 15 |
| | Partial list of genes up-regulated in the kidney of GLIS2 [GeneID=84662] knockout mice compared to the wild type. | 101 | 2.04502e-316 | 6.81134e-315 | 2.1987 | 2375 | 7550 | 1.73% | 1 | 83 |
| | Genes up-regulated in primary keratinocytes by expression of constantly active NOTCH1 [GeneID=4851]. | 101 | 2.04502e-316 | 6.81134e-315 | 2.1987 | 2375 | 7550 | 1.73% | 1 | 29 |
| | Genes up-regulated in MEF cells (embryonic fibroblast) lacking TP53 and BRCA1 [GeneID=7157;672] by expression of TP53; most genes are further up-regulated by simultaneous expression of BRCA1. | 101 | 2.04502e-316 | 6.81134e-315 | 2.1987 | 2375 | 7550 | 1.73% | 1 | 37 |
| | Genes down-regulated in erythroid lineage cells by RUNX1-RUNX1T1 [GeneID=861;862] fusion . | 101 | 2.04502e-316 | 6.81134e-315 | 2.1987 | 2375 | 7550 | 1.73% | 1 | 15 |
| | Proteins secreted in co-culture of LKR-13 tumor cells (non-small cell lung cancer, NSCLC) and MEC stroma cells (endothelium). | 101 | 2.04502e-316 | 6.81134e-315 | 2.1987 | 2375 | 7550 | 1.73% | 1 | 66 |
| | Genes down-regulated in CD133+ [GeneID=8842] cells (hematopoietic stem cells, HSC) compared to the CD133- cells. | 114 | 1.23948e-287 | 3.65754e-286 | 2.1932 | 2172 | 6922 | 1.58% | 4 | 217 |
| | Genes up-regulated in PC-3 cells (prostate cancer) stably expressing ST7 [GeneID=7982] off a plasmid vector. | 120 | 1.26305e-271 | 3.54074e-270 | 2.1538 | 2141 | 6948 | 1.56% | 1 | 85 |
| | Genes from the turquoise module which are down-regulated in HAEC cells (primary aortic endothelium) after exposure to the oxidized 1-palmitoyl-2-arachidonyl-sn-3-glycerophosphocholine (oxPAPC). | 134 | 5.70447e-246 | 1.43208e-244 | 2.1527 | 1941 | 6302 | 1.42% | 2 | 53 |
| | Cluster 3 of aberrantly hypomethylated genes in blasts from AML (acute myeloid leukemia) patients. | 138 | 1.58430e-240 | 3.86202e-239 | 2.0744 | 2079 | 7005 | 1.52% | 1 | 42 |
| | Genes down-regulated in primary fibroblast cell culture at 30 min time point after infection with HCMV (AD169 strain). | 139 | 4.90097e-238 | 1.18610e-236 | 2.0444 | 2135 | 7299 | 1.56% | 4 | 146 |
| | Genes up-regulated in NSCLC (non-small cell lung carcinoma) cell lines resistant to gefitinib [PubChem=123631] compared to the sensitive ones. | 146 | 2.00399e-223 | 4.61742e-222 | 2.4011 | 1380 | 4017 | 1.01% | 1 | 80 |
| | Genes down-regulated in ER(-) / PR(-) breast tumors (do not express ESR1 and PGR [GeneID=2099;5241]) with molecular similarity to ER(+) (class A) relative to the rest of the ER(-) / PR(-) samples (class B). | 146 | 2.00399e-223 | 4.61742e-222 | 2.4011 | 1380 | 4017 | 1.01% | 1 | 33 |
| | Genes up-regulated in MDA-MB-231 cells (breast cancer) upon overexpression of PARVB [GeneID=29780] under all three culture conditions. | 146 | 2.00399e-223 | 4.61742e-222 | 2.4011 | 1380 | 4017 | 1.01% | 1 | 6 |
| | Genes down-regulated in glioma cell lines treated with both decitabine [PubChem=451668] and TSA [PubChem=5562]. | 146 | 2.00399e-223 | 4.61742e-222 | 2.4011 | 1380 | 4017 | 1.01% | 1 | 12 |
| | Up-regulated 'cooperation response genes': responded synergystically to the combination of mutant TP53 and HRAS [GeneID=7157;3265] in YAMC cells (colon). | 146 | 2.00399e-223 | 4.61742e-222 | 2.4011 | 1380 | 4017 | 1.01% | 1 | 25 |
| | Genes enriched in oligodendrocytes in the adult mouse brain identified through correlation-based searches seeded with the oligodendrocyte cell-type specific gene expression patterns. | 151 | 1.65403e-220 | 3.68487e-219 | 2.0251 | 2027 | 6996 | 1.48% | 3 | 74 |
| | Genes regulated in MCF7 cells (breast cancer) by expression of the full-length form of ERBB2 [GeneID=2064] at 60 h time point. | 186 | 8.67604e-174 | 1.56915e-172 | 2.4040 | 1069 | 3108 | 0.78% | 2 | 27 |
| | Hepatic graft versus host disease (GVHD), day 7: down-regulated in allogeneic vs syngeneic bone marrow transplant. | 188 | 3.57173e-172 | 6.39112e-171 | 3.0204 | 691 | 1599 | 0.50% | 1 | 41 |
| | Genes with intermediate-CpG-density promoters (ICP) bearing histone H3 trimethylation mark at K4 (H3K4me3) in brain. | 199 | 8.06614e-171 | 1.36354e-169 | 2.0776 | 1469 | 4942 | 1.07% | 1 | 29 |
| | Genes with intermediate-CpG-density promoters (ICP) bearing histone H3 trimethylation mark at K4 (H3K4me3) in ES cells (embryonic stem). | 199 | 8.06614e-171 | 1.36354e-169 | 2.0776 | 1469 | 4942 | 1.07% | 1 | 30 |
| | Genes down-regulated in partially reprogrammed and pluripotent cell populations (induced, iPS; and embryonic stem cells, ES) compared to parental lineage-commited cell lines. | 199 | 8.06614e-171 | 1.36354e-169 | 2.0776 | 1469 | 4942 | 1.07% | 1 | 7 |
| | Genes from the grey module which are up-regulated in HAEC cells (primary aortic endothelium) after exposure to the oxidized 1-palmitoyl-2-arachidonyl-sn-3-glycerophosphorylcholine (oxPAPC). | 238 | 1.11432e-124 | 1.57503e-123 | 2.1881 | 942 | 3009 | 0.69% | 1 | 16 |
| | Down-regulated genes in colon carcinoma tumors compared to the matched normal mucosa samples. | 238 | 1.11432e-124 | 1.57503e-123 | 2.1881 | 942 | 3009 | 0.69% | 1 | 31 |
| | Selected genes implicated in metastasis and epithelial-to-mesenchymal transition (EMT) which were up-regulated in MDA-MB-231 cells (breast cancer) upon knockdown of SKI [GeneID=6497] by RNAi. | 238 | 1.11432e-124 | 1.57503e-123 | 2.1881 | 942 | 3009 | 0.69% | 1 | 17 |
| | Down-regulated in HEK293 cells (kidney epithelium) by treatment with sodium arsenite [PubChem=26435]. | 238 | 1.11432e-124 | 1.57503e-123 | 2.1881 | 942 | 3009 | 0.69% | 1 | 17 |
| | The optimal set of 70 prognostic markers predicting poor breast cancer clinical outcome (defined as developing metastases with 5 years). | 262 | 5.17364e-111 | 6.64280e-110 | 2.0901 | 936 | 3130 | 0.68% | 1 | 55 |
| | Top 20 genes whose down-regulation correlated with gastrointestinal stromal tumors (GIST) and synovial sarcoma compared to other tumors. | 264 | 3.83129e-110 | 4.88200e-109 | 2.1756 | 842 | 2705 | 0.61% | 2 | 20 |
| | UV only responding genes in primary fibroblasts from young donors. | 266 | 2.32534e-108 | 2.94077e-107 | 2.3524 | 694 | 2062 | 0.51% | 1 | 61 |
| | Genes commonly down-regulated in CD-1 and CD-2 clusters of multiple myeloma samples and which were higher expressed in the CD-1 group. | 268 | 1.26193e-105 | 1.58400e-104 | 2.1894 | 795 | 2538 | 0.58% | 2 | 49 |
| | Cluster 2 of aberrantly hypomethylated genes in blasts from AML (acute myeloid leukemia) patients. | 276 | 2.46485e-97 | 3.00426e-96 | 2.4399 | 576 | 1650 | 0.42% | 1 | 8 |
| | Genes down-regulated in MDA-MB-231 cells (breast cancer, mutated TP53 [GeneID=7157]) undergoing mitotic arrest and apoptosis after treatment with 100 nM docetaxel [PubChem=148124]. | 276 | 2.46485e-97 | 3.00426e-96 | 2.4399 | 576 | 1650 | 0.42% | 1 | 18 |
| | Granule constituents expressed during mouse promyelocytic cell line differentiation to neutrophils. | 276 | 2.46485e-97 | 3.00426e-96 | 2.4399 | 576 | 1650 | 0.42% | 1 | 25 |
| | Top 20 negative significant genes associated with synovial sarcoma tumors. | 276 | 2.46485e-97 | 3.00426e-96 | 2.4399 | 576 | 1650 | 0.42% | 1 | 20 |
| | Genes down-regulated in glioma cell lines after knockdown of SPARC [GeneID=6678] by RNAi. | 276 | 2.46485e-97 | 3.00426e-96 | 2.4399 | 576 | 1650 | 0.42% | 1 | 14 |
| | Genes down-regulated in the mouse lung cancer model and which reverted to normal levels upon treatment with bexarotene [PubChem=82146]. | 276 | 2.46485e-97 | 3.00426e-96 | 2.4399 | 576 | 1650 | 0.42% | 1 | 28 |
| | Genes silenced by DNA methylation in bladder cancer cell lines. | 276 | 2.46485e-97 | 3.00426e-96 | 2.4399 | 576 | 1650 | 0.42% | 1 | 53 |
| | Genes whose expression most strongly and consistently associated with the short term survival of patients with high grade glioma tumors. | 276 | 2.46485e-97 | 3.00426e-96 | 2.4399 | 576 | 1650 | 0.42% | 1 | 9 |
| | Genes most significantly down-regulated in multiple myeloma samples, compared to normal bone marrow plasma cells. | 276 | 2.46485e-97 | 3.00426e-96 | 2.4399 | 576 | 1650 | 0.42% | 1 | 40 |
| | Genes up-regulated in HeLa cells (cervical carcinoma) at 4 h after stimulation with TNF [GeneID=7124]. | 276 | 2.46485e-97 | 3.00426e-96 | 2.4399 | 576 | 1650 | 0.42% | 1 | 54 |
| | Downregulated in the gastrocnemius muscle of aged adult mice (30-month) vs young adult (5-month) | 317 | 4.71219e-79 | 5.00057e-78 | 2.2312 | 566 | 1773 | 0.41% | 1 | 49 |

| | |
|---|---|
| The test set contains 137,142 (14%) of all 958,541 regions. | The test set picked 595 genes, the background set picked 600 genes. |
| *MSigDB Perturbation* has 3,364 terms covering 17,091 (95%) of all 18,041 genes. | 3,364 ontology terms were tested (100%) using an annotation count range of [1, Inf]. |
| GREAT version 3.0.0 | |
| Species assembly: hg19 | |
| Association rule: Basal+extension: 5000 bp upstream, 1000 bp downstream, 1000000 bp max extension, curated regulatory domains included | |

Figure S10: Part 2 out of 2 of GREAT results table for the MSigDB Perturbations ontology for results from RADMeth analysis.

#### S2.3 Validation of the LuxUS preanalysis step

To validate the F-test in the preanalysis step, we compared the Bayes factor values for the 9945 genomic windows that had preanalysis F-test p-value $\leq 0.1$ to the Bayes factor values for 10000 randomly chosen genomic windows which had preanalysis F-test p-value $> 0.1$. The same coverage requirements were applied to the both sets of genomic windows. In Fig. S11 the cumulative proportion of Bayes factors is plotted as the function of Bayes factor value, which has been plotted in logarithmic scale. The proportion of small Bayes factors is considerably higher amongst the genomic windows with preanalysis F-test p-value$>$ 0.1 than for the genomic windows which had the preanalysis F-test p-value $\leq 0.1$. We conclude that using the preanalysis step leads to discarding only a few genomic windows with high Bayes factor values.

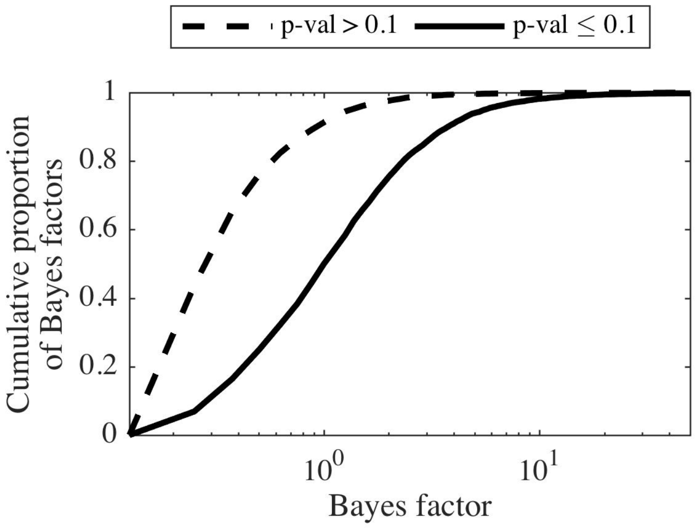


Figure S11: Cumulative proportion of Bayes factors as a function of Bayes factor value for the colon cancer data set. Logarithmic scale is used for the x-axis. In this figure the cumulative proportions are shown up to Bayes factor value 50. The dashed line represents the Bayes factors for the randomly picked 10000 genomic windows that had p-value $> 0.1$ and the solid line represents the genomic windows with p-value $\leq 0.1$ in the preanalysis test. The figure shows that the proportion of small Bayes factors is greater for the cytosines with greater p-values in the preanalysis phase.

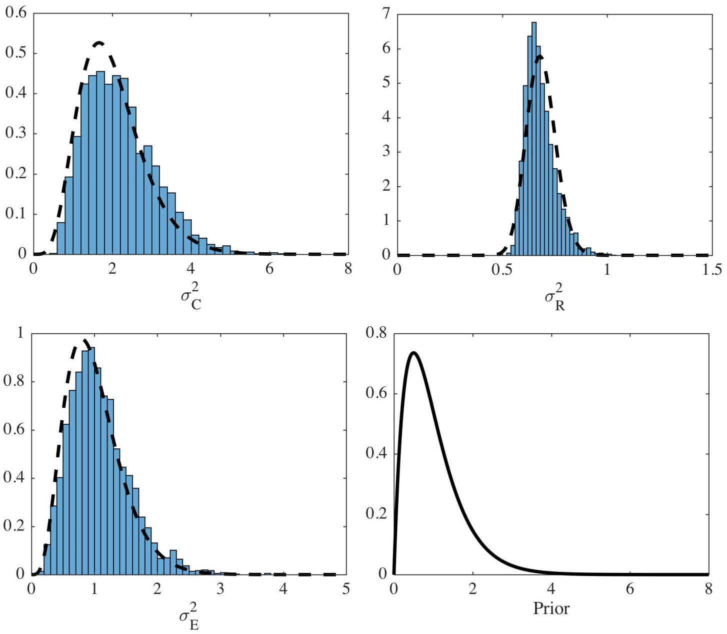


Figure S12: Histograms of the posterior sample means for the variance parameters $\sigma^2_C$, $\sigma^2_R$ and $\sigma^2_E$ from the colon cancer WGBS-seq data analysis and the probability density function of their prior, Gamma(2,2). The figure shows how much the posterior distributions for the parameters have diverged from the prior distributions. The results from this real data analysis also indicate that the magnitudes of these variance terms are different. The probability density functions for the estimated gamma distributions for the variance parameters have been plotted with dashed line on top of the histograms. The estimated gamma distributions are used as priors for the respective parameters in the analysis of the simulated data.

#### S2.4 Setting simulation parameters based on the colon cancer WGBS data set

The analysis of the colon cancer WGBS-seq data was also used to set certain parameters for the simulation experiments to realistic values. Gamma(2,2) prior was used for the variance parameters $\sigma^2_C$, $\sigma^2_R$ and $\sigma^2_E$ when estimating the model for the colon cancer data. The means of the posterior samples for these parameters across all windows were stored and their distributions can be seen in Fig. S12. Gamma densities were fitted to the sample means using gamfit function from Matlab. For this purpose only the genomic windows with 10 or more cytosines and with minimum mean sample coverage of 5 were considered. The resulting shape and scale parameters for the gamma distributions were approximately $(6, 1/3)$, $(98, 1/143)$ and $(5, 1/5)$, respectively for $\sigma^2_C$, $\sigma^2_R$

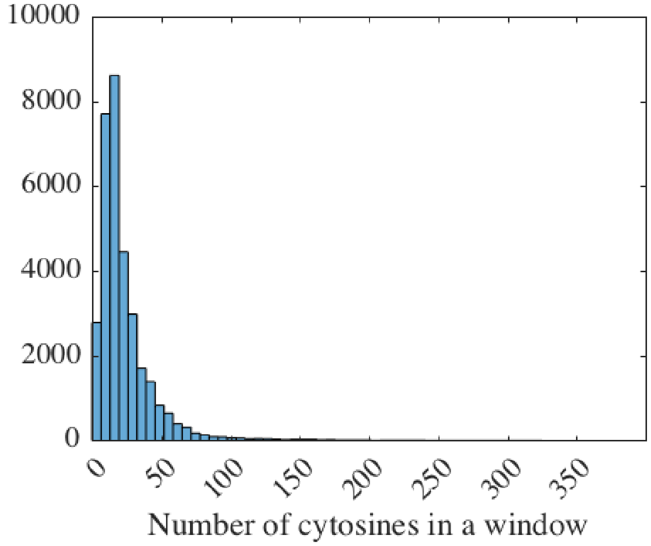


Figure S13: Histograms of the number of cytosines in genomic windows of length of 2000 basepairs.

and $\sigma_E^2$.

The average number of cytosines in a genomic window for the colon cancer data set was also explored. The windows of 2000 bp were first formed with only filtering the cytosines so that there must be at least one sample present from both case and control groups for a cytosine to be included in a genomic window. This resulted in 16210 windows in chromosome 21 and 16502 windows in chromosome 22, and the distribution of number of cytosines in these windows is shown in Fig. S13. The mean of the number of cytosines in the 2000 basepair windows is 22.8 and the mode is 12. Based on this result, we decided to use cytosine frequency of 10 cytosines in a 1000 basepair window for the simulation experiments.

### S3 Analysis of lung cancer RRBS-seq data set

#### S3.1 Estimating variance parameters

To compare the estimated variance parameters for the colon cancer data set to another real bisulfite sequencing data set, we ran LuxUS to non-small cell lung cancer (NSCLC) data set by Hascher et al. (2014). The data is available with GEO accession number GSE52140. We chose eight of the samples in the data set: the normal and highly metastatic cell line samples for both cell lines

A549 and HTB56. We left the samples where treatment had been applied out of the analysis. The design matrix included the intercept term, cell line (A549 or HTB56) indicator variable and indicator variable for highly metastatic cell line status.

The data is available as preprocessed count type data. We ran the LuxUS preanalysis to the count data and applied the same restrictions as for the colon cancer data set. Namely, the maximum number of cytosines in a genomic window was 20, maximum length of the window was 2000bp and the F-test p-value threshold for the highly metastatic cell line covariate was 0.1. Each cytosine that was added to a genomic window had to have at least one sample from both normal cell line and highly metastatic cell line groups with coverage of 5. Also, there had to be at least one sample with mean coverage over 5 from both cell line groups for the genomic window to be accepted. Out of the genomic windows that passed the preanalysis phase, we chose randomly 500 windows with 10 or more cytosines in them from five largest chromosomes for LuxUS analysis. There were less than 500 of such windows in chromosome 5, so the windows for chromosome 6 were used instead.

The means of the posterior samples for the three variance parameters $\sigma_C^2$, $\sigma_R^2$ and $\sigma_E^2$ were stored for every genomic window. Chromosome-wise boxplots of the estimates can be found from Fig. S14-S16. From the boxplots we can see, that there is variation between the individual genomic windows, which can be seen as the variance of the posterior means. The variance parameter $\sigma_R^2$ has the narrowest range, while the range for $\sigma_C^2$ is the widest. The similar shapes of the boxplots indicate that the distribution of the posterior sample means are still very similar between different chromosomes. Also, the posterior means for the three variance parameters are different from each other, even though the same prior was used for all of them. This suggests that these parameters are not very sensitive to the choice of the hyperprior parameters.

If the boxplots of the posterior means for the NSCLC data set are compared to the histograms of the posterior sample means for the colon cancer data set presented in Fig. S12, we can see that the ranges of the distributions and the relative order of the widths of the ranges for the three variance parameters are very similar. So based on the distribution of the posterior means for a number of windows the sources of variation are similar for these two data sets, but the posterior mean for a single genomic window has room for variation.

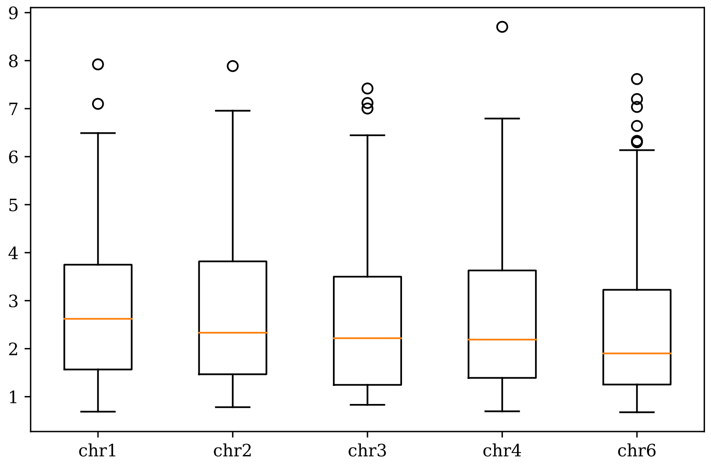


Figure S14: Boxplots of the $\sigma_C^2$ posterior sample means for 500 windows for five chromosomes.

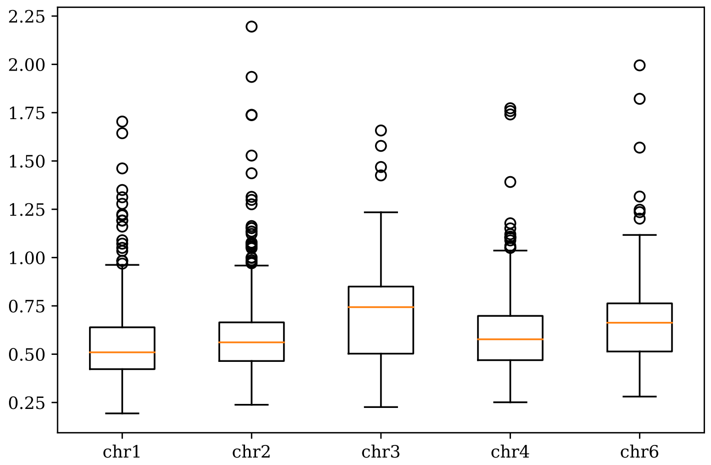


Figure S15: Boxplots of the $\sigma_R^2$ posterior sample means for 500 windows for five chromosomes.

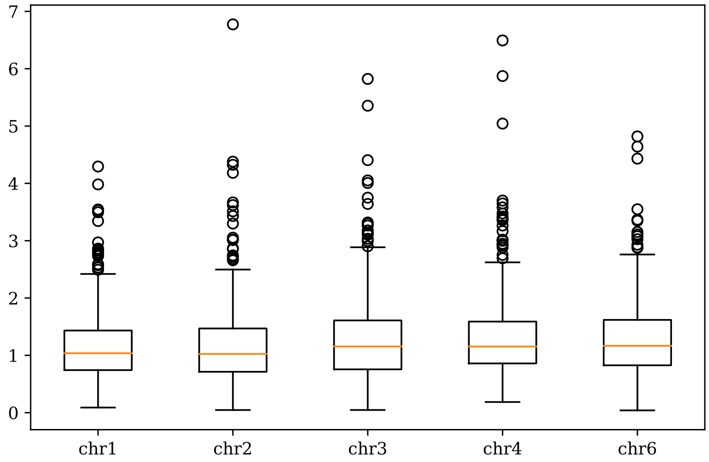


Figure S16: Boxplots of the $\sigma_E^2$ posterior sample means for 500 windows for five chromosomes.

#### S3.2 Sensitivity to $\sigma_b^2$ value

To see how changing the $\sigma_b^2$ value affects the estimated Bayes factors, and the performance as a consequence, we ran LuxUS analysis with three different $\sigma_b^2$ values and compared the resulting Bayes factors. We used the lung cancer data set by Hascher et al. (2014) for the comparison. The same preprocessing as in Supplementary information Section S3.1 was used. In Section S3.1 the $\sigma_b^2$ was set to 15. In here we additionally tested setting $\sigma_b^2$ to values 5 and 25, to see how this would affect the Bayes factor values. We ran the LuxUS analysis with the different $\sigma_b^2$ values for the same randomly picked 500 genomic windows from chromosome 1 with at least 10 cytosines in each as in Section S3.1. Overall, the BFs obtained with different $\sigma_b^2$ values are quite similar. In here we compare the Bayes factors calculated with $\sigma_b^2 = 15$ to the ones calculated with $\sigma_b^2 = 5$ and $\sigma_b^2 = 25$. The Pearson correlation coefficients were 1.000 for both comparisons. The Supplementary Fig. S17-S18 show the comparison as scatter plots. Most of the points lie close to the $y = x$ line which was drawn for comparison, indicating that analysis with $\sigma_b^2$ set to 15 or 5 and 15 or 25 would give similar Bayes factor values for the 500 genomic windows. An inherent feature of the Savage-Dickey density ratio estimator is that a smaller value of the variance parameter results in a higher Bayes factor estimate as can be seen from our results in Supplementary Figs. S17-S18. Based on this comparison it seems that LuxUS is not very sensitive to the $\sigma_b^2$ value.

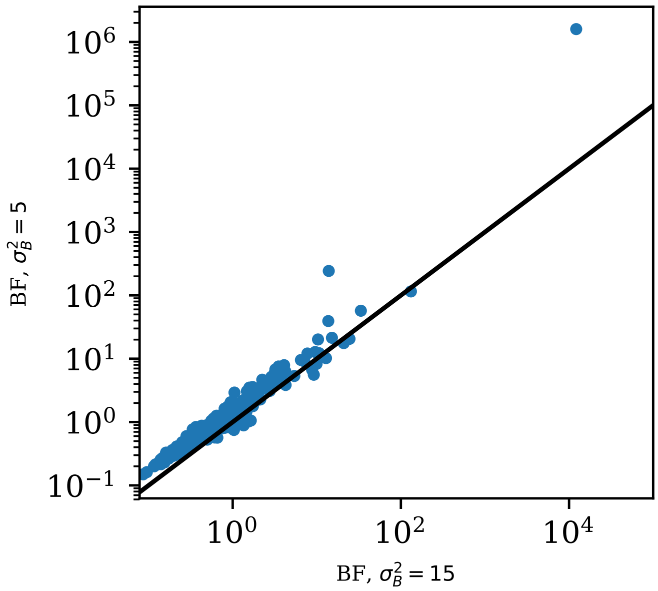


Figure S17: Bayes factors calculated with different $\sigma_b^2$ values plotted against each other, with both x- and y-axis log-scaled. The values in the x-axis have been calculated with $\sigma_b^2 = 15$ and the values in the y-axis with $\sigma_b^2 = 5$. The black line shows line for $y = x$ for comparison.

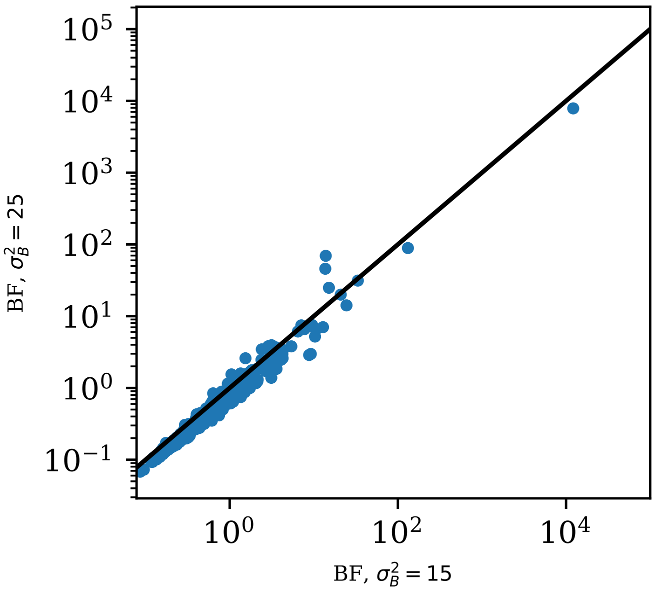


Figure S18: Bayes factors calculated with different $\sigma_b^2$ values plotted against each other, with both x- and y-axis log-scaled. The values in the x-axis have been calculated with $\sigma_b^2 = 15$ and the values in the y-axis with $\sigma_b^2 = 25$. The black line shows line for $y = x$ for comparison.

### S4 Performance comparisons on simulated BS-seq data

#### S4.1 BS-seq data simulation

The simulation of BS-seq data was conducted in the following way. First, 100 sets of true values of the experimental parameters $\text{bs}_{\text{eff}}$, $\text{bs}^*_{\text{eff}}$ and $\text{seq}_{\text{err}}$ were generated from distributions Beta$(99, 1)$, Beta$(1, 999)$ and Beta$(1, 999)$ respectively. For $\text{seq}_{\text{err}}$ and $\text{bs}^*_{\text{eff}}$ we use the means of their respective true distributions as inputs when estimating the model. For $\text{bs}_{\text{eff}}$, we simulate the usage of spike-in control cytosines from lambda phage genome. For each parameter set, we draw 3066 random numbers from binomial distribution, one for each cytosine in the lambda phage genome. For each draw the number of trials is 10 to represent sequencing coverage of 10, and the success rate is the true value of $\text{bs}_{\text{eff}}$ for the respective data set. The bisulfite conversion rate estimates for each parameter set are then calculated as the ratio of sum of total number generated methylated reads per the total number of reads for the whole lambda phage genome. The genomic coordinates are also generated randomly. The coordinates are retrieved from range $[1, 1000]$ using random integer generation function provided in NumPy. The number of cytosines for each data set is set to 10. After this, for each of the 100 sets of generated experimental parameters and genomic coordinates we generate two data sets, one with differential methylation and one without differential methylation, totaling in 200 data sets. The proportion of differentially methylated regions (DMRs) is 50% in this case. Additionally, we generated more simulated data sets without differential methylation to prepare a simulation setup with a smaller and more realistic DMR proportion. We generated 950 data sets without differential methylation in total. When combined with 50 of the data sets with differential methylation, the total data set count was 1000 and the corresponding DMR proportion was 5%.

To generate a BS-seq data set with the simulated experimental parameters, the true $\beta$ and the design matrix are set first. The number of replicates in the simulated experiments was varied and the used values were 6, 12 and 24. $\beta_0$ is generated from normal distribution with mean $\mu_{B,0}$ and variance $\sigma_B^2 = 0.25$, while $\beta_1$ is set to $\mu_{B,1}$ if there is differential methylation and to zero if there is no differential methylation. The used $\boldsymbol{\mu}_B = (\mu_{B,0}, \mu_{B,1})$ values were $(-1.4, 1)$, $(-1.4, 2.3)$, $(1.4, -1)$ and $(1.4, -2.3)$, which correspond to approximate values of 0.2, 0.5, 0.2 and 0.5 of $\Delta\theta$, the difference in methylation proportions between the case and control groups, respectively. The methylation states corresponding to the $\mu_{B,0} = -1.4$ and $\mu_{B,0} = 1.4$ are approximately 0.2 and 0.8 respectively, representing realistic methylation levels often observed in bisulfite sequencing experiments for cell populations. During the inference, the variance $\sigma_B^2$ is set to 15. The design matrix includes column for the intercept term and column indicating whether the replicate is a case or a control. Only binary variables were used in the simulation experiments, as the RADMeth tool to which we compare LuxUS does not support continuous variables. Also, two of the other

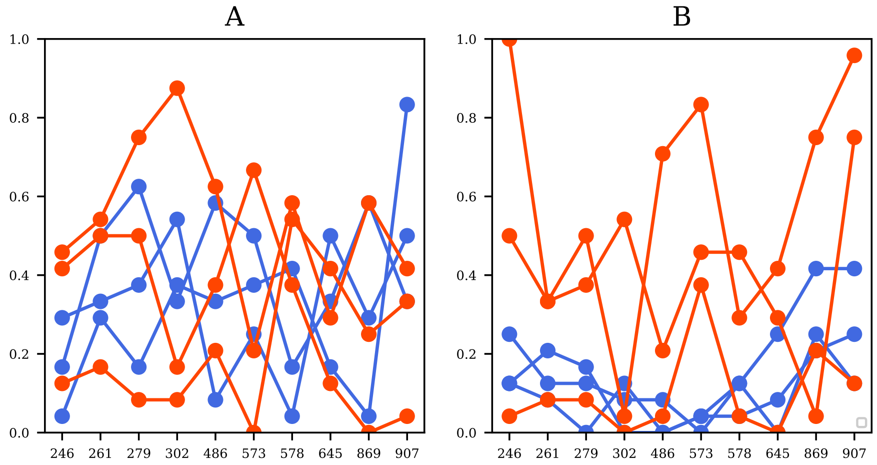


Figure S19: Two examples of simulated data sets. The methylation proportion $N_{BS,C}/N_{BS}$ (y-axis) is plotted with dots for each cytosine with their genomic coordinates in the x-axis. The dots for the same replicate are connected with a line. Controls are plotted with blue and cases with red. The simulated genomic coordinates and experimental parameters are the same for both figures A and B. The number of replicates $N_R$ is 6 and the read coverage $N_{BS}$ is 24. In A, $\mu_b = (1.4, 0)$ was used for simulation, so there is no differential methylation between the cases and controls and the replicates seem to be mixed. In B, $\mu_b = (1.4, -1)$ was used to induce differential methylation between case and control replicates. Using this $\mu_b$ value the expected difference in methylation states between the cases and controls is $|\Delta\theta| \approx 0.2$ We can see that there is differential methylation in B as most of the cases plotted in red seem to be separated from the blue controls.

tools we compare our method to, metilene and bsseq, only support comparison of two groups.

To demonstrate the advantages of taking confounding covariates into account and allowing a general experimental design in the analysis, we also generated data with a more complex experimental design. In this setting, the design matrix consisted of columns for an intercept term, a binary covariate distinguishing cases from control samples, a confounding binary covariate and a confounding continuous covariate. The case-control binary covariate and the confounding binary covariate were balanced in the design matrix. With 12 replicates the design matrix for this setting would be of form

$$\mathbf{D} = \begin{bmatrix} 1 & 0 & 0 & r_1 \\ 1 & 1 & 0 & r_2 \\ 1 & 0 & 1 & r_3 \\ 1 & 1 & 1 & r_4 \\ 1 & 0 & 0 & r_5 \\ 1 & 1 & 0 & r_6 \\ 1 & 0 & 1 & r_7 \\ 1 & 1 & 1 & r_8 \\ 1 & 0 & 0 & r_9 \\ 1 & 1 & 0 & r_{10} \\ 1 & 0 & 1 & r_{11} \\ 1 & 1 & 1 & r_{12} \end{bmatrix}. \quad (3)$$

The values for the confounding continuous covariate, $r_i$, $i = 1, ..., 12$ , were set to random numbers from range $[0, 1]$. The values for the number of replicates used in the simulations were 12 and 24. The used $\boldsymbol{\mu}_B = (\mu_{B,0}, \mu_{B,1}, \mu_{B,2}, \mu_{B,3})$ values were $(1.4, -1, 2, -3)$. Again, $\beta_0$ is generated from a normal distribution with mean $\mu_{B,0}$ and variance $\sigma_B^2 = 0.25$, while $\beta_1$, $\beta_2$ and $\beta_3$ are set to the corresponding $\mu_{B,i}$, $i = 1, 2, 3$ values given in $\boldsymbol{\mu}_B$. This simulated data set can be found as a zipped file from the LuxUS GitHub repository.

The random effects for replicates and cytosines are generated from normal distributions with zero mean and variances $\sigma_R^2$ and $\sigma_C^2$. Then $Y$ is generated from normal distribution with the sum of the fixed linear term and the random effects as the mean and $\sigma_E$ as the variance. The variances $\sigma_C^2$, $\sigma_R^2$ and $\sigma_E^2$ were set to the means of the gamma distributions that were fitted using the posterior samples for the Hansen data set. Then $\theta$ is calculated from $Y$, and $p_{\text{BS},C}$ is calculated using $\theta$ and the experimental parameters. Finally, the number of methylated reads is generated from binomial distribution with success rate $p_{\text{BS},C}$ and varying the number of reads. The values used for the number of reads were 6, 12 and 24. Two examples of data sets simulated with the described method are presented in Supplementary Fig. S19.

The data is then saved as Stan input files. The priors for $\sigma_C^2$, $\sigma_R^2$ and $\sigma_E^2$ were set using the results from the real data analysis and $\sigma_B^2$ is set to 15 for the model estimation. The data was also saved into formats that are supported by RADMeth, M$^3$D, dmrseq, DSS, metilene and bsseq tools. The preanalysis step with F-test was not performed for the simulated data, as the aim was to compute the Bayes factors for all simulated genomic windows for the purposes of calculating receiver operating characteristic ROC curve. Each of the genomic windows, i.e. data sets, is labeled as differentially methylated or not differentially methylated. Each of the simulated genomic windows were analysed separately. In the ROC calculation for LuxUS, each genomic window is considered as one data point. The number of genomic windows with differential methylation is exactly half of the total number of windows, allowing balanced ROC calculation. For the other tools which provide a cytosine-specific p-value or Bayes factor, ROC calculation was done in cytosine-wise manner. For this purpose the label of the genomic window (differentially methylated or not differentially methylated) was given to all the cytosines in the window. For the simulation setups with 5% DMR proportion the analysis was carried out similarly with each of the tools being compared. To present the results for these simulation setups, we chose to use precision-recall curves, which are commonly used when the proportions of the positive and negative classes are unbalanced. The precision-recall curve demonstrates the tradeoff between precision, i.e. the number of true positives over the sum of number of true and false positives, and recall, i.e. the number of true positives over the sum of number of true positivies and number of false negatives, for different threshold values. The results can be summarised by calculating area under precision-recall curves or average precision (AP), defined as

$$AP = \sum_n (R_n - R_{n-1})P_n, \tag{4}$$

in Scikit-learn Python package (Pedregosa et al., 2011) whose implementation we use. $R_n$ and $P_n$ are recall and precision for threshold $n$.

#### S4.2 Computation times for LuxUS

Table S1: Comparison of mean computation times in seconds for estimating the LuxUS model with HMC and ADVI for a single genomic window, using simulated data set with $\boldsymbol{\mu}_b = (1.4, -1)$. $N_{\mathrm{BS}}$ denotes the number of sequencing reads overlapping a cytosine and $N_R$ denotes the number of samples.

| $N_{\mathrm{BS}}$ | $N_R$ | HMC | ADVI |
|---|---|---|---|
| 6 | 6 | 84.136 | 3.489 |
| 6 | 12 | 503.055 | 13.178 |
| 6 | 24 | 2918.109 | 55.816 |
| 12 | 6 | 83.792 | 3.193 |
| 12 | 12 | 448.537 | 11.249 |
| 12 | 24 | 2671.042 | 53.291 |
| 24 | 6 | 78.075 | 2.961 |
| 24 | 12 | 400.321 | 10.306 |
| 24 | 24 | 2504.799 | 48.830 |

#### S4.3 ROC curves

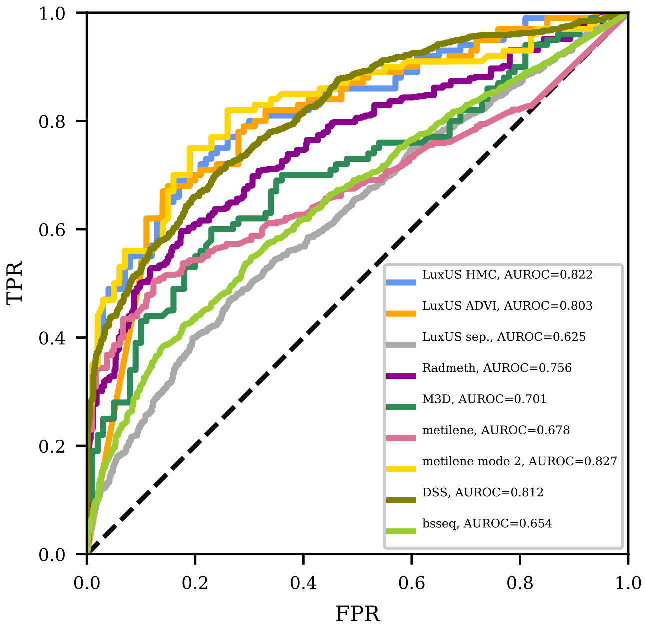


Figure S20: Receiver operating characteristics curves for LuxUS with HMC and ADVI model estimation, LuxUS separately for each cytosine, RADMeth, M$^3$D, DSS, metilene and bsseq. The dashed black line shows the expected ROC curve for random guessing. Results for simulated data with $\mu_B = (1.4, -1)$, 12 reads and 12 replicates was used for the figure.

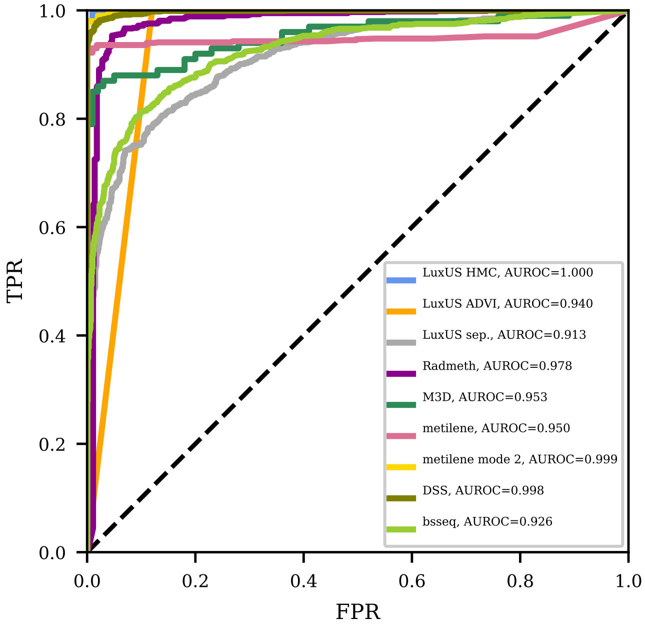


Figure S21: Receiver operating characteristics curves for LuxUS with HMC and ADVI model estimation, LuxUS separately for each cytosine, RADMeth, M$^3$D, DSS, metilene and bsseq. The dashed black line shows the expected ROC curve for random guessing. Results for simulated data with $\mu_B = (1.4, -2.3)$, 12 reads and 12 replicates was used for the figure.

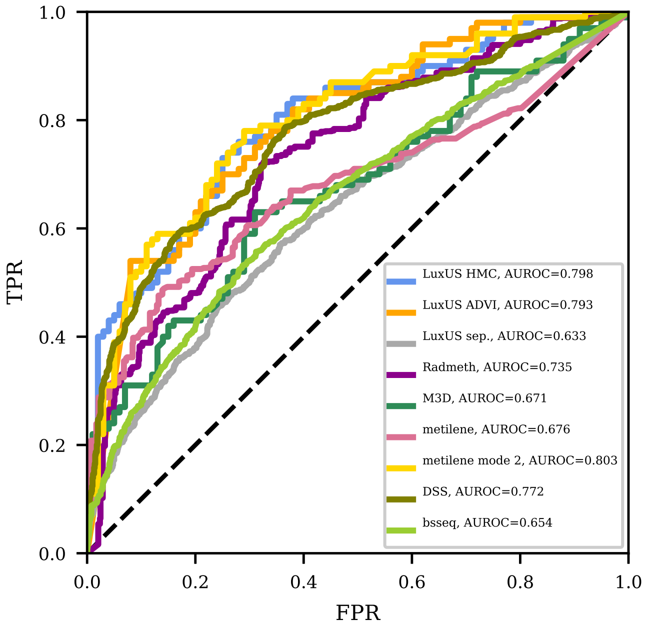


Figure S22: Receiver operating characteristics curves for LuxUS with HMC and ADVI model estimation, LuxUS separately for each cytosine, RADMeth, $M^3D$, DSS, metilene and bsseq. The dashed black line shows the expected ROC curve for random guessing. Results for simulated data with $\mu_B = (-1.4, 1)$, 12 reads and 12 replicates was used for the figure.

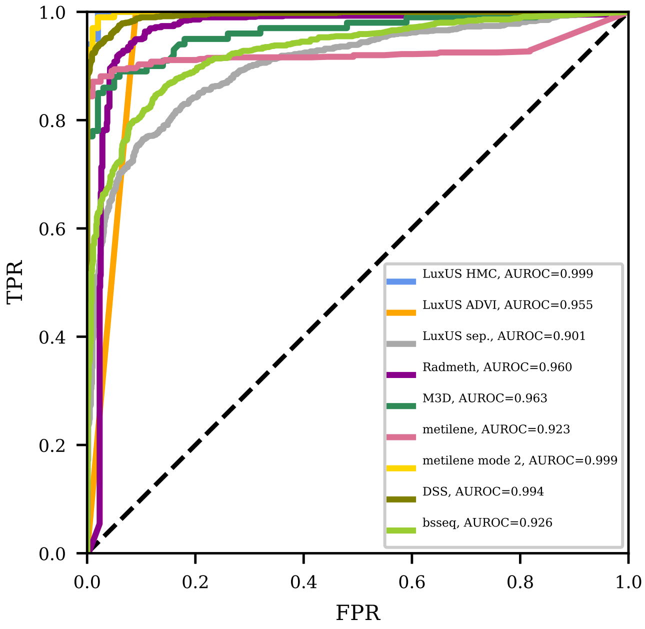


Figure S23: Receiver operating characteristics curves for LuxUS with HMC and ADVI model estimation, LuxUS separately for each cytosine, RADMeth, M$^3$D, DSS, metilene and bsseq. The dashed black line shows the expected ROC curve for random guessing. Results for simulated data with $\mu_B = (-1.4, 2.3)$, 12 reads and 12 replicates was used for the figure.

#### S4.4 AUROC value tables

Table S2: AUROC values for LuxUS with HMC and VI, LuxUS for separate cytosines, RADMeth, $M^3D$, DSS, metilene and bsseq for simulated data set with $\mu_B = (-1.4, 2.3)$, corresponding to $\Delta\theta \approx 0.5$. The highest AUROC value is bolded for each case. $N_{\mathrm{BS}}$ denotes the number of sequencing reads overlapping a cytosine and $N_R$ denotes the number of samples.

| $N_{\mathrm{BS}}$ | $N_R$ | LuxUS HMC | LuxUS ADVI | LuxUS sep. | RAD-Meth | $M^3D$ | metilene | metilene mode 2 | DSS | bsseq |
|---|---|---|---|---|---|---|---|---|---|---|
| 6 | 6 | 0.932 | 0.881 | 0.740 | 0.836 | 0.905 | 0.815 | **0.936** | 0.929 | 0.790 |
| 6 | 12 | 0.996 | 0.910 | 0.851 | 0.936 | 0.963 | 0.915 | **0.997** | 0.993 | 0.894 |
| 6 | 24 | **1.000** | 0.945 | 0.964 | 0.983 | 0.952 | 0.969 | 0.998 | 0.998 | 0.968 |
| 12 | 6 | **0.966** | 0.931 | 0.793 | 0.895 | 0.907 | 0.804 | 0.962 | 0.950 | 0.811 |
| 12 | 12 | **0.999** | 0.955 | 0.901 | 0.960 | 0.963 | 0.923 | **0.999** | 0.994 | 0.926 |
| 12 | 24 | **1.000** | 0.935 | 0.981 | 0.993 | 0.966 | 0.974 | **1.000** | **1.000** | 0.980 |
| 24 | 6 | **0.975** | 0.956 | 0.816 | 0.929 | 0.909 | 0.868 | 0.972 | 0.961 | 0.830 |
| 24 | 12 | **1.000** | 0.965 | 0.939 | 0.978 | 0.942 | 0.935 | 0.999 | 0.995 | 0.945 |
| 24 | 24 | **1.000** | 0.940 | 0.993 | 0.999 | 0.965 | 0.973 | **1.000** | **1.000** | 0.989 |

Table S3: AUROC values for LuxUS with HMC and ADVI, LuxUS for separate cytosines, RADMeth, $M^3D$, DSS, metilene and bsseq for simulated data set with $\mu_B = (-1.4, 1)$, corresponding to $\Delta\theta \approx 0.2$. The highest AUROC value is bolded for each case. $N_{\mathrm{BS}}$ denotes the number of sequencing reads overlapping a cytosine and $N_R$ denotes the number of samples.

| $N_{\mathrm{BS}}$ | $N_R$ | LuxUS HMC | LuxUS ADVI | LuxUS sep. | RAD-Meth | $M^3D$ | metilene | metilene mode 2 | DSS | bsseq |
|---|---|---|---|---|---|---|---|---|---|---|
| 6 | 6 | 0.716 | 0.711 | 0.557 | 0.599 | 0.691 | 0.583 | **0.729** | 0.718 | 0.603 |
| 6 | 12 | 0.847 | 0.839 | 0.633 | 0.752 | 0.667 | 0.664 | **0.848** | 0.845 | 0.658 |
| 6 | 24 | **0.946** | 0.908 | 0.741 | 0.849 | 0.716 | 0.773 | 0.927 | 0.915 | 0.744 |
| 12 | 6 | 0.660 | 0.665 | 0.516 | 0.630 | 0.670 | 0.624 | **0.682** | 0.676 | 0.586 |
| 12 | 12 | 0.798 | 0.793 | 0.633 | 0.735 | 0.671 | 0.676 | **0.803** | 0.772 | 0.654 |
| 12 | 24 | 0.924 | 0.902 | 0.727 | 0.873 | 0.757 | 0.827 | **0.931** | 0.909 | 0.754 |
| 24 | 6 | 0.713 | 0.687 | 0.573 | 0.634 | 0.702 | 0.581 | **0.725** | 0.691 | 0.615 |
| 24 | 12 | 0.824 | 0.807 | 0.663 | 0.763 | 0.656 | 0.725 | **0.836** | 0.809 | 0.673 |
| 24 | 24 | 0.952 | 0.939 | 0.790 | 0.917 | 0.731 | 0.815 | **0.954** | 0.933 | 0.802 |

#### S4.5 Precision-recall curves

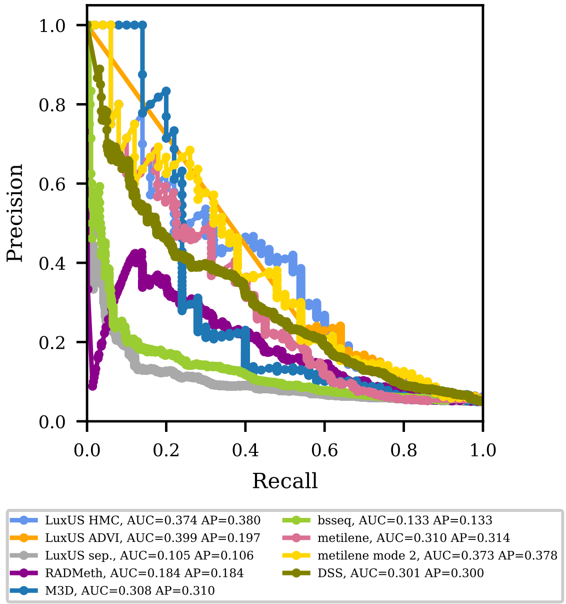


Figure S24: Precision-recall curves for LuxUS with HMC and ADVI model estimation, LuxUS separately for each cytosine, RADMeth, M$^3$D, DSS, metilene and bsseq. Results for simulated data with $\mu_B = (1.4, -1)$, 12 reads and 12 replicates was used for the figure. The curve has been plotted based on 1000 simulated data sets, where the proportion of differentially methylated regions was 5%. The area under the precision-recall curve (AUC) and average precision (AP) for each method are given in the legend.

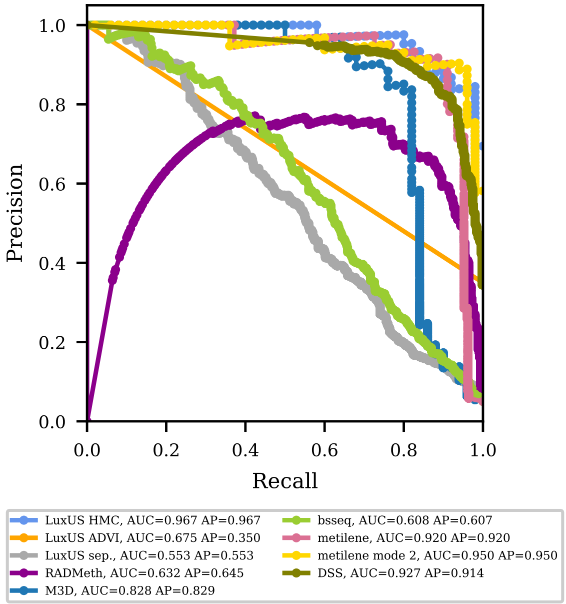


Figure S25: Precision-recall curves for LuxUS with HMC and ADVI model estimation, LuxUS separately for each cytosine, RADMeth, M$^3$D, DSS, metilene and bsseq. Results for simulated data with $\mu_B = (1.4, -2.3)$, 12 reads and 12 replicates was used for the figure. The curve has been plotted based on 1000 simulated data sets, where the proportion of differentially methylated regions was 5%. The area under the precision-recall curve (AUC) and average precision (AP) for each method are given in the legend.

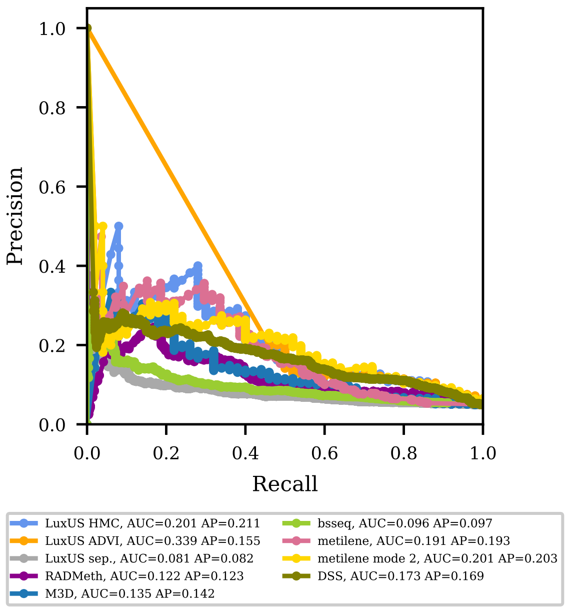


Figure S26: Precision-recall curves for LuxUS with HMC and ADVI model estimation, LuxUS separately for each cytosine, RADMeth, M$^3$D, DSS, metilene and bsseq. Results for simulated data with $\mu_B = (-1.4, 1)$, 12 reads and 12 replicates was used for the figure. The curve has been plotted based on 1000 simulated data sets, where the proportion of differentially methylated regions was 5%. The area under the precision-recall curve (AUC) and average precision (AP) for each method are given in the legend.

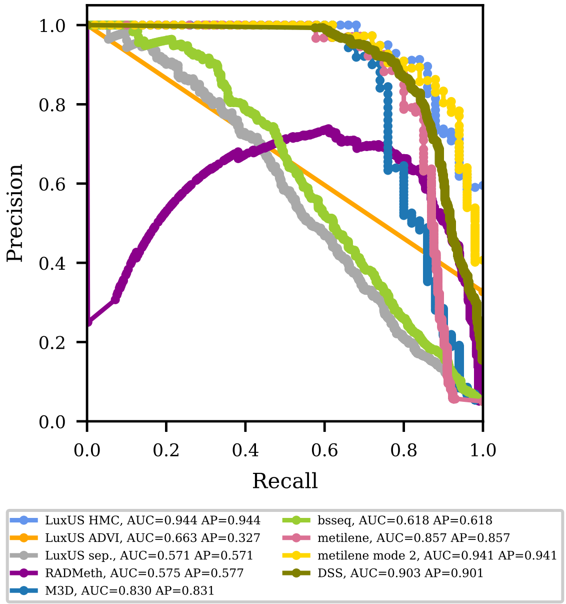


Figure S27: Precision-recall curves for LuxUS with HMC and ADVI model estimation, LuxUS separately for each cytosine, RADMeth, M$^3$D, DSS, metilene and bsseq. Results for simulated data with $\mu_B = (-1.4, 2.3)$, 12 reads and 12 replicates was used for the figure. The curve has been plotted based on 1000 simulated data sets, where the proportion of differentially methylated regions was 5%. The area under the precision-recall curve (AUC) and average precision (AP) for each method are given in the legend.

#### S4.6 Average precision tables

Table S4: Average precision values for LuxUS with HMC and ADVI, LuxUS for separate cytosines, RADMeth, $M^3D$, DSS, metilene and bsseq for simulated data set with $\mu_B = (1.4, -1)$, corresponding to $\Delta\theta \approx -0.2$. The highest AP value is bolded for each case. These values have been calculated based on 1000 simulated data sets, where the proportion of differentially methylated regions was 5%. $N_{\mathrm{BS}}$ denotes the number of sequencing reads overlapping a cytosine and $N_R$ denotes the number of samples.

| $N_{\mathrm{BS}}$ | $N_R$ | LuxUS HMC | LuxUS ADVI | LuxUS sep. | RAD-Meth | $M^3D$ | metilene | metilene mode 2 | DSS | bsseq |
|---|---|---|---|---|---|---|---|---|---|---|
| 6 | 6 | 0.127 | 0.105 | 0.059 | 0.094 | 0.123 | 0.121 | **0.162** | 0.143 | 0.073 |
| 6 | 12 | **0.279** | 0.161 | 0.096 | 0.163 | 0.158 | 0.195 | 0.275 | 0.248 | 0.110 |
| 6 | 24 | **0.652** | 0.343 | 0.169 | 0.307 | 0.252 | 0.472 | 0.606 | 0.576 | 0.211 |
| 12 | 6 | 0.184 | 0.120 | 0.068 | 0.101 | 0.165 | 0.137 | **0.226** | 0.198 | 0.087 |
| 12 | 12 | **0.380** | 0.197 | 0.106 | 0.184 | 0.310 | 0.314 | 0.378 | 0.300 | 0.133 |
| 12 | 24 | 0.631 | 0.295 | 0.197 | 0.309 | 0.242 | 0.549 | **0.635** | 0.577 | 0.233 |
| 24 | 6 | 0.153 | 0.095 | 0.070 | 0.081 | 0.164 | 0.184 | **0.187** | 0.155 | 0.087 |
| 24 | 12 | 0.415 | 0.203 | 0.131 | 0.245 | 0.252 | 0.407 | **0.463** | 0.394 | 0.163 |
| 24 | 24 | **0.717** | 0.266 | 0.269 | 0.431 | 0.311 | 0.567 | 0.655 | 0.602 | 0.262 |

Table S5: Average precision values for LuxUS with HMC and ADVI, LuxUS for separate cytosines, RADMeth, $M^3D$, DSS, metilene and bsseq for simulated data set with $\mu_B = (1.4, -2.3)$, corresponding to $\Delta\theta \approx -0.5$. The highest AP value is bolded for each case. These values have been calculated based on 1000 simulated data sets, where the proportion of differentially methylated regions was 5%. $N_{\mathrm{BS}}$ denotes the number of sequencing reads overlapping a cytosine and $N_R$ denotes the number of samples.

| $N_{\mathrm{BS}}$ | $N_R$ | LuxUS HMC | LuxUS ADVI | LuxUS sep. | RAD-Meth | $M^3D$ | metilene | metilene mode 2 | DSS | bsseq |
|---|---|---|---|---|---|---|---|---|---|---|
| 6 | 6 | **0.776** | 0.323 | 0.246 | 0.360 | 0.634 | 0.630 | 0.772 | 0.682 | 0.259 |
| 6 | 12 | **0.964** | 0.365 | 0.483 | 0.596 | 0.846 | 0.851 | 0.959 | 0.933 | 0.552 |
| 6 | 24 | **1.000** | 0.347 | 0.841 | 0.767 | 0.913 | 0.915 | 0.998 | 0.990 | 0.860 |
| 12 | 6 | **0.718** | 0.337 | 0.267 | 0.349 | 0.639 | 0.581 | 0.698 | 0.647 | 0.327 |
| 12 | 12 | **0.967** | 0.350 | 0.553 | 0.645 | 0.829 | 0.920 | 0.950 | 0.914 | 0.607 |
| 12 | 24 | **0.998** | 0.331 | 0.875 | 0.788 | 0.884 | 0.956 | 0.997 | 0.980 | 0.879 |
| 24 | 6 | **0.753** | 0.329 | 0.329 | 0.389 | 0.583 | 0.635 | 0.731 | 0.645 | 0.359 |
| 24 | 12 | **0.958** | 0.321 | 0.656 | 0.578 | 0.857 | 0.903 | 0.944 | 0.906 | 0.685 |
| 24 | 24 | **1.000** | 0.321 | 0.934 | 0.781 | 0.963 | 0.987 | **1.000** | 0.997 | 0.931 |

Table S6: Average precision values for LuxUS with HMC and ADVI, LuxUS for separate cytosines, RADMeth, M$^3$D, DSS, metilene and bsseq for simulated data set with $\mu_B = (-1.4, 1)$, corresponding to $\Delta\theta \approx 0.2$. The highest AP value is bolded for each case. These values have been calculated based on 1000 simulated data sets, where the proportion of differentially methylated regions was 5%. $N_{\mathrm{BS}}$ denotes the number of sequencing reads overlapping a cytosine and $N_R$ denotes the number of samples.

| $N_{\mathrm{BS}}$ | $N_R$ | LuxUS HMC | LuxUS ADVI | LuxUS sep. | RAD-Meth | M$^3$D | metilene | metilene mode 2 | DSS | bsseq |
|---|---|---|---|---|---|---|---|---|---|---|
| 6 | 6 | 0.147 | 0.119 | 0.064 | 0.104 | 0.124 | 0.128 | **0.186** | 0.155 | 0.075 |
| 6 | 12 | **0.364** | 0.185 | 0.095 | 0.175 | 0.245 | 0.212 | 0.348 | 0.344 | 0.114 |
| 6 | 24 | **0.666** | 0.307 | 0.208 | 0.310 | 0.293 | 0.512 | 0.655 | 0.588 | 0.246 |
| 12 | 6 | **0.132** | 0.119 | 0.061 | 0.095 | 0.094 | 0.105 | 0.122 | 0.122 | 0.073 |
| 12 | 12 | **0.211** | 0.155 | 0.082 | 0.123 | 0.142 | 0.193 | 0.203 | 0.169 | 0.097 |
| 12 | 24 | **0.637** | 0.273 | 0.236 | 0.339 | 0.328 | 0.559 | 0.626 | 0.537 | 0.268 |
| 24 | 6 | **0.260** | 0.123 | 0.097 | 0.127 | 0.219 | 0.155 | 0.213 | 0.192 | 0.110 |
| 24 | 12 | 0.364 | 0.168 | 0.139 | 0.185 | 0.269 | 0.365 | **0.409** | 0.365 | 0.159 |
| 24 | 24 | 0.702 | 0.303 | 0.307 | 0.481 | 0.332 | 0.639 | **0.729** | 0.664 | 0.323 |

Table S7: Average precision values for LuxUS with HMC and ADVI, LuxUS for separate cytosines, RADMeth, M$^3$D, DSS, metilene and bsseq for simulated data set with $\mu_B = (-1.4, 2.3)$, corresponding to $\Delta\theta \approx 0.5$. The highest AP value is bolded for each case. These values have been calculated based on 1000 simulated data sets, where the proportion of differentially methylated regions was 5%. $N_{\mathrm{BS}}$ denotes the number of sequencing reads overlapping a cytosine and $N_R$ denotes the number of samples.

| $N_{\mathrm{BS}}$ | $N_R$ | LuxUS HMC | LuxUS ADVI | LuxUS sep. | RAD-Meth | M$^3$D | metilene | metilene mode 2 | DSS | bsseq |
|---|---|---|---|---|---|---|---|---|---|---|
| 6 | 6 | 0.754 | 0.304 | 0.274 | 0.337 | 0.685 | 0.646 | **0.763** | 0.733 | 0.317 |
| 6 | 12 | **0.958** | 0.350 | 0.442 | 0.541 | 0.818 | 0.821 | 0.957 | 0.919 | 0.498 |
| 6 | 24 | **0.997** | 0.370 | 0.786 | 0.674 | 0.894 | 0.968 | 0.985 | 0.986 | 0.817 |
| 12 | 6 | **0.829** | 0.337 | 0.332 | 0.384 | 0.747 | 0.653 | 0.791 | 0.778 | 0.379 |
| 12 | 12 | **0.944** | 0.327 | 0.571 | 0.577 | 0.831 | 0.857 | 0.941 | 0.901 | 0.618 |
| 12 | 24 | **0.999** | 0.298 | 0.867 | 0.702 | 0.932 | 0.982 | 0.998 | 0.978 | 0.879 |
| 24 | 6 | **0.688** | 0.276 | 0.269 | 0.380 | 0.616 | 0.578 | 0.667 | 0.540 | 0.301 |
| 24 | 12 | 0.989 | 0.352 | 0.688 | 0.657 | 0.855 | 0.943 | **0.993** | 0.931 | 0.721 |
| 24 | 24 | **1.000** | 0.298 | 0.926 | 0.775 | 0.910 | 0.977 | **1.000** | 0.991 | 0.916 |

#### S4.7 Estimating the methylation proportions $\boldsymbol{\theta}$

In Fig. S28 the boxplots of the posterior means of $\boldsymbol{\theta}$ from the LuxUS runs are plotted, combining the means for all cytosines and replicates for a simulation scenario where there is no differential methylation. The true methylation state without noise terms was 0.8 and the number of replicates was 6. The number of

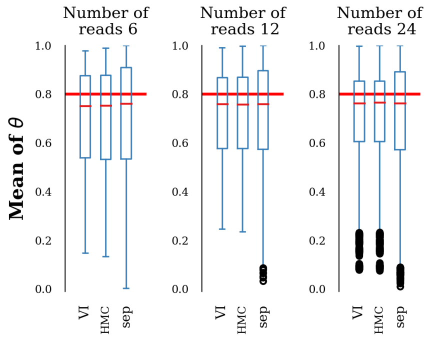


Figure S28: Boxplots of the estimated $\theta$ parameter posterior means for the 200 simulated data sets, where number of replicates was 6 and $\boldsymbol{\mu}_b = (1.4, -1)$ was used for the simulations. The number of reads was 6, 12 and 24 respectively in the figures from left to right. The thick red lines show the true methylation level.

reads was varied to see whether increasing the number of reads would increase the precision of the estimates. The boxplots are plotted for the LuxUS model, estimated with both HMC and ADVI methods, and for the separate cytosine analysis, where model estimation was done with HMC sampling. The boxplots are wider for the separate cytosine analysis than for the whole window analysis, indicating higher variance in the estimates. There is no clear difference between the HMC and ADVI approaches for the whole window analysis. Increasing the number of reads from 6 to 12 and to 24 decreases the variation of the estimates, but the shift from 12 to 24 reads seems to increase the number of outliers.

To conclude, these simulation experiment results show that the variance of the methylation state estimates is smaller if a larger genomic window is analysed at the same time. The means of the samples for the methylation states $\boldsymbol{\theta}$ are approximately as close to the true value in both cases.

#### S4.8 ROC curves for simulation setting with two confounding covariates

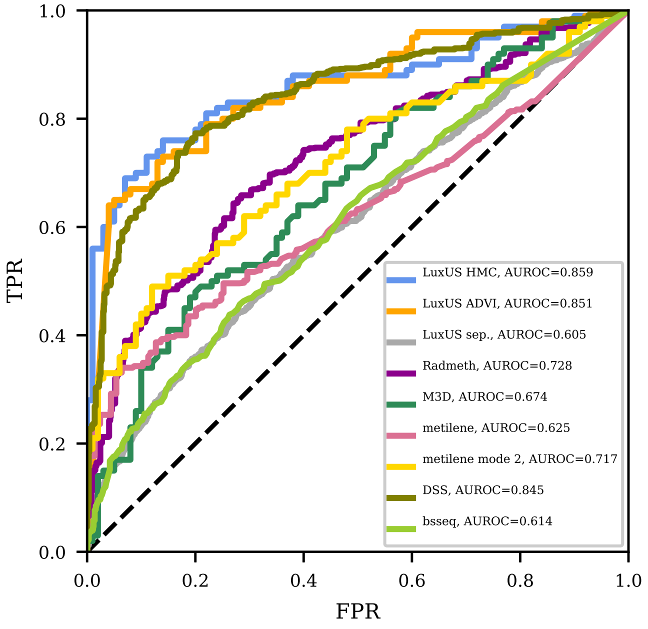


Figure S29: Receiver operating characteristics curves for LuxUS with HMC and ADVI model estimation, LuxUS separately for each cytosine, RADMeth, $M^3D$, DSS, metilene and bsseq. Results for simulated data with confounding covariates, 6 reads and 12 replicates were used for the figure. The curves have been plotted based on 200 simulated data sets, where the proportion of differentially methylated regions was 50%. The AUROC values are given in the legend.

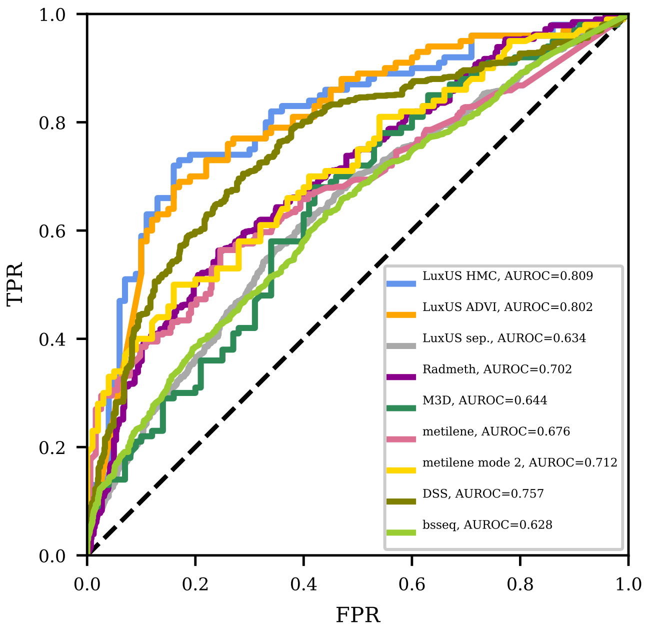


Figure S30: Receiver operating characteristics curves for LuxUS with HMC and ADVI model estimation, LuxUS separately for each cytosine, RADMeth, $M^3D$, DSS, metilene and bsseq. Results for simulated data with confounding covariates, 12 reads and 12 replicates were used for the figure. The curves have been plotted based on 200 simulated data sets, where the proportion of differentially methylated regions was 50%. The AUROC values are given in the legend.

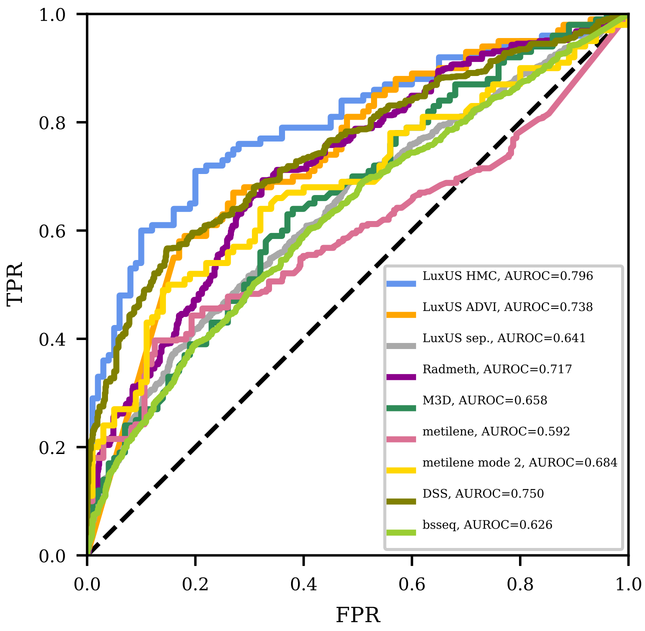


Figure S31: Receiver operating characteristics curves for LuxUS with HMC and ADVI model estimation, LuxUS separately for each cytosine, RADMeth, M$^3$D, DSS, metilene and bsseq. Results for simulated data with confounding covariates, 24 reads and 12 replicates were used for the figure. The curves have been plotted based on 200 simulated data sets, where the proportion of differentially methylated regions was 50%. The AUROC values are given in the legend.

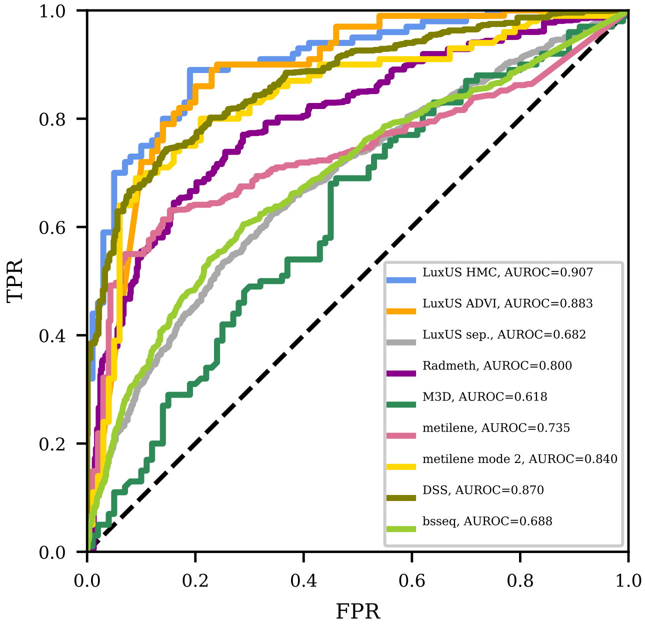


Figure S32: Receiver operating characteristics curves for LuxUS with HMC and ADVI model estimation, LuxUS separately for each cytosine, RADMeth, M$^3$D, DSS, metilene and bsseq. Results for simulated data with confounding covariates, 6 reads and 24 replicates was used for the figure. The curves have been plotted based on 200 simulated data sets, where the proportion of differentially methylated regions was 50%. The AUROC values are given in the legend.

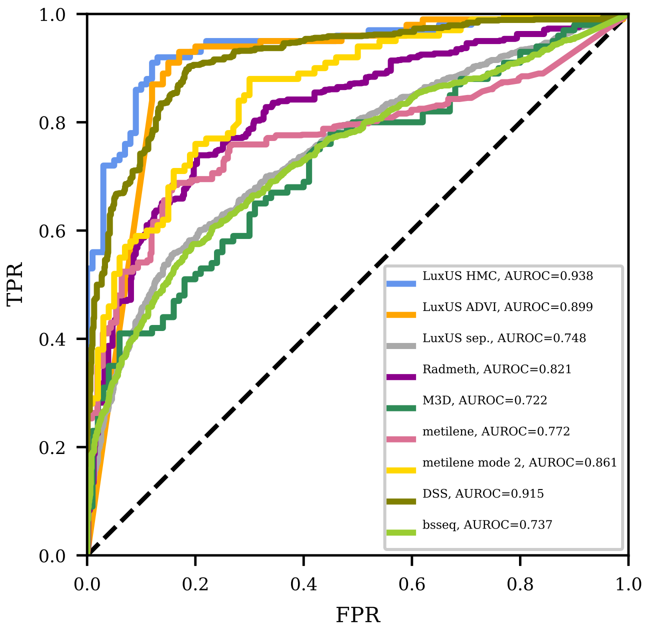


Figure S33: Receiver operating characteristics curves for LuxUS with HMC and ADVI model estimation, LuxUS separately for each cytosine, RADMeth, $M^3D$, DSS, metilene and bsseq. Results for simulated data with confounding covariates, 12 reads and 24 replicates were used for the figure. The curves have been plotted based on 200 simulated data sets, where the proportion of differentially methylated regions was 50%. The AUROC values are given in the legend.

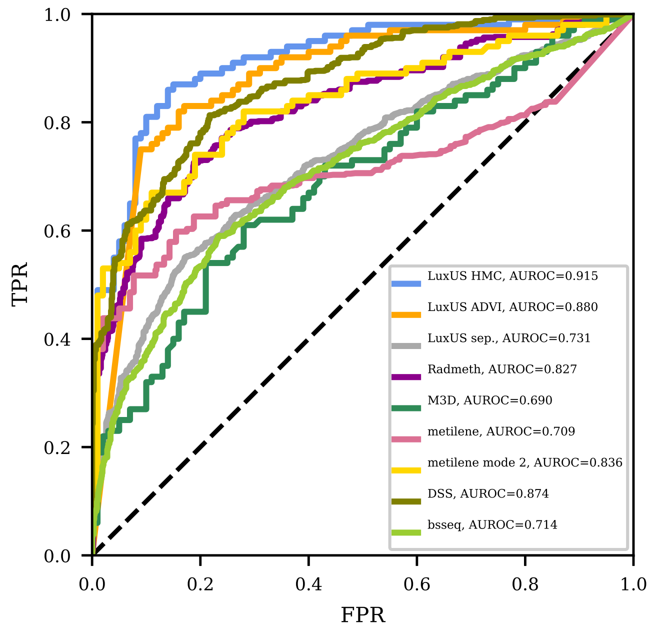


Figure S34: Receiver operating characteristics curves for LuxUS with HMC and ADVI model estimation, LuxUS separately for each cytosine, RADMeth, $M^3D$, DSS, metilene and bsseq. Results for simulated data with confounding covariates, 24 reads and 24 replicates were used for the figure. The curves have been plotted based on 200 simulated data sets, where the proportion of differentially methylated regions was 50%. The AUROC values are given in the legend.

#### S4.9 Precision-recall curves for simulation setting with two confounding covariates

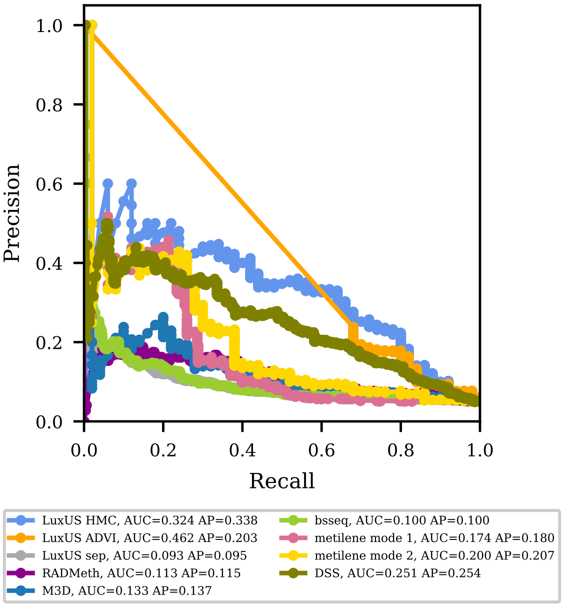


Figure S35: Precision-recall curves for LuxUS with HMC and ADVI model estimation, LuxUS separately for each cytosine, RADMeth, M$^3$D, DSS, metilene and bsseq. Results for simulated data with confounding covariates, 6 reads and 12 replicates were used for the figure. The curves have been plotted based on 1000 simulated data sets, where the proportion of differentially methylated regions was 5%. The area under the precision-recall curve (AUC) and average precision (AP) for each method are given in the legend.

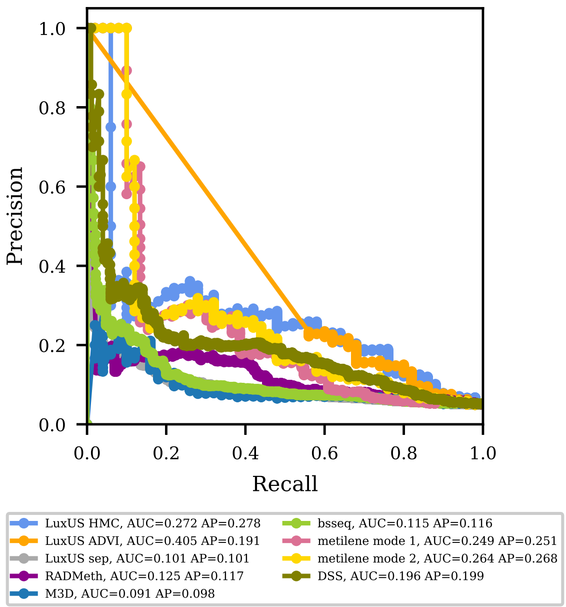


Figure S36: Precision-recall curves for LuxUS with HMC and ADVI model estimation, LuxUS separately for each cytosine, RADMeth, M$^3$D, DSS, metilene and bsseq. Results for simulated data with confounding covariates, 12 reads and 12 replicates were used for the figure. The curves have been plotted based on 1000 simulated data sets, where the proportion of differentially methylated regions was 5%. The area under the precision-recall curve (AUC) and average precision (AP) for each method are given in the legend.

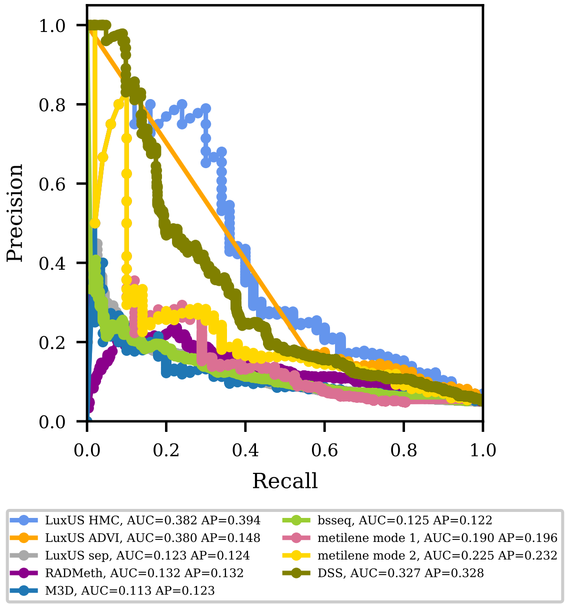


Figure S37: Precision-recall curves for LuxUS with HMC and ADVI model estimation, LuxUS separately for each cytosine, RADMeth, M$^3$D, DSS, metilene and bsseq. Results for simulated data with confounding covariates, 24 reads and 12 replicates were used for the figure. The curves have been plotted based on 1000 simulated data sets, where the proportion of differentially methylated regions was 5%. The area under the precision-recall curve (AUC) and average precision (AP) for each method are given in the legend.

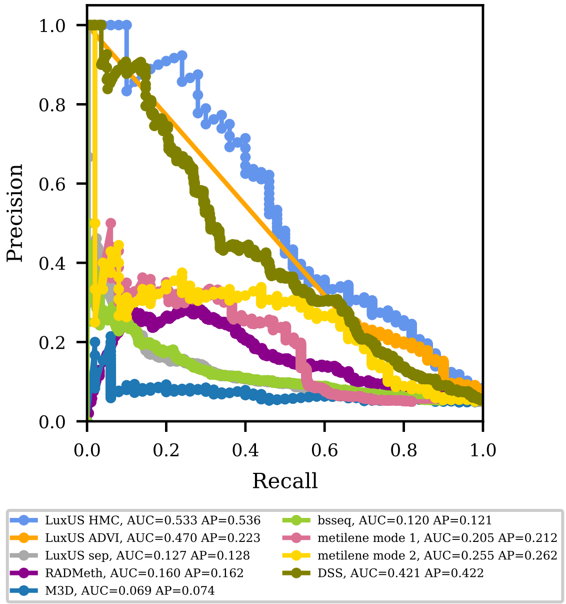


Figure S38: Precision-recall curves for LuxUS with HMC and ADVI model estimation, LuxUS separately for each cytosine, RADMeth, M$^3$D, DSS, metilene and bsseq. Results for simulated data with confounding covariates, 6 reads and 24 replicates were used for the figure. The curves have been plotted based on 1000 simulated data sets, where the proportion of differentially methylated regions was 5%. The area under the precision-recall curve (AUC) and average precision (AP) for each method are given in the legend.

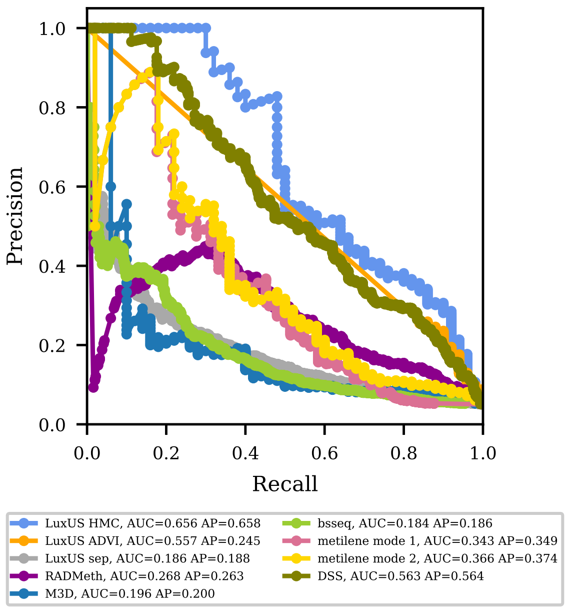


Figure S39: Precision-recall curves for LuxUS with HMC and ADVI model estimation, LuxUS separately for each cytosine, RADMeth, M$^3$D, DSS, metilene and bsseq. Results for simulated data with confounding covariates, 12 reads and 24 replicates were used for the figure. The curves have been plotted based on 1000 simulated data sets, where the proportion of differentially methylated regions was 5%. The area under the precision-recall curve (AUC) and average precision (AP) for each method are given in the legend.

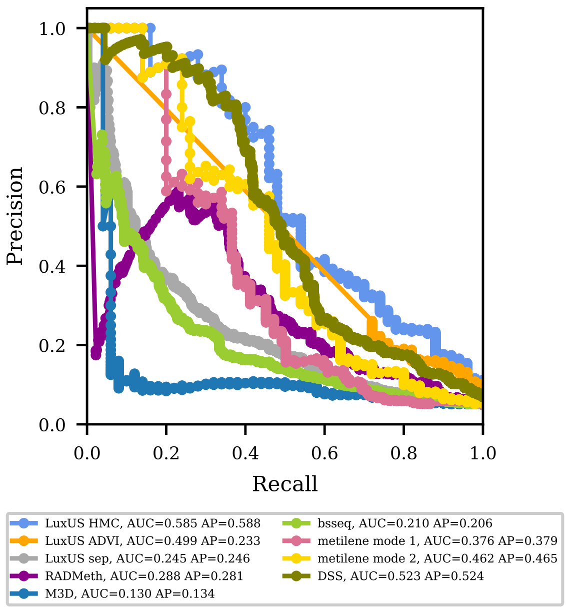


Figure S40: Precision-recall curves for LuxUS with HMC and ADVI model estimation, LuxUS separately for each cytosine, RADMeth, M$^3$D, DSS, metilene and bsseq. Results for simulated data with confounding covariates, 24 reads and 24 replicates were used for the figure. The curves have been plotted based on 1000 simulated data sets, where the proportion of differentially methylated regions was 5%. The area under the precision-recall curve (AUC) and average precision (AP) for each method are given in the legend.

#### S4.10 Average precision table for simulation setting with two confounding covariates

Table S8: Average precision values for LuxUS with HMC and ADVI, LuxUS for separate cytosines, RADMeth, $M^3D$, DSS, metilene and bsseq for simulated data set with confounding covariates. The highest AP value is bolded for each case. These values have been calculated based on 1000 simulated data sets, where the proportion of differentially methylated regions was 5%. $N_{\mathrm{BS}}$ denotes the number of sequencing reads overlapping a cytosine and $N_R$ denotes the number of samples.

| $N_{\mathrm{BS}}$ | $N_R$ | LuxUS HMC | LuxUS ADVI | LuxUS sep. | RAD-Meth | $M^3D$ | metilene | metilene mode 2 | DSS | bsseq |
|---|---|---|---|---|---|---|---|---|---|---|
| 6 | 12 | **0.338** | 0.203 | 0.095 | 0.115 | 0.137 | 0.180 | 0.207 | 0.254 | 0.100 |
| 6 | 24 | **0.536** | 0.223 | 0.128 | 0.162 | 0.074 | 0.212 | 0.262 | 0.422 | 0.121 |
| 12 | 12 | **0.278** | 0.191 | 0.101 | 0.117 | 0.098 | 0.251 | 0.268 | 0.199 | 0.116 |
| 12 | 24 | **0.658** | 0.245 | 0.188 | 0.263 | 0.200 | 0.349 | 0.374 | 0.564 | 0.186 |
| 24 | 12 | **0.394** | 0.148 | 0.124 | 0.132 | 0.123 | 0.196 | 0.232 | 0.328 | 0.122 |
| 24 | 24 | **0.588** | 0.233 | 0.246 | 0.281 | 0.134 | 0.379 | 0.465 | 0.524 | 0.206 |

### S5 Performance comparisons on simulated differentially methylated regions based on real BS-seq data

#### S5.1 Preanalysis and setting the priors

Additional performance comparisons were made with simulated data set which is based on RRBS-seq data of 12 control samples. Six of the controls were turned to cases by adding 10000 simulated differentially methylated regions to randomly chosen CpG islands. The size of the DMRs and the methylation difference between the cases and controls was varied. For this comparison the CpG islands containing a DMR were divided into DMR and non-DMR sets. For both sets the LuxUS preanalysis step was run to divide them into genomic windows, with maximum width of 2000 bp and maximum number of cytosines of 20. The genomic windows were defined so that all tools could analyse them without computational problems. This meant filtering out windows with only one cytosine, filtering out cytosines for which either all control or all case samples had zero coverage and filtering out cytosines for which the methylation state for both the case and control samples was either 0 or 1. The latter of the restrictions was applied because of computational problems in RADMeth tool, even if it would remove some of the cytosines which could contain valuable information for other tools. No other coverage restrictions were applied to the genomic windows. Also, there was no preanalysis F-test restrictions for the genomic windows. As the experimental parameters were not known for the original data set, we set $\text{BS}_{\text{eff}} = 1$, $\text{BS}^*_{\text{eff}} = 0$ and $\text{seq}_{\text{err}} = 0$. For LuxUS analysis, the hyperprior parameters and value for $\sigma^2_C$ were set in the same way as for the data simulated from LuxUS model.

### References


Alp Kucukelbir, Rajesh Ranganath, Andrew Gelman, and David M. Blei. Automatic variational inference in stan, 2015.

David M. Blei, Alp Kucukelbir, and Jon D. McAuliffe. Variational inference: A review for statisticians. *Journal of the American Statistical Association*, 112(518):859–877, 2017. doi: 10.1080/01621459.2017.1285773. URL https://doi.org/10.1080/01621459.2017.1285773.

Andrew Gelman, Donald Rubin, David Dunson, Hal S. Stern, Aki Vehtari, and John Carlin. *Bayesian Data Analysis, Third Edition*. CRC Press, Boca Raton, Florida, 2014.

Antje Hascher, Ann-Kristin Haase, Katja Hebestreit, Christian Rohde, Hans-Ulrich Klein, Maria Rius, Dominik Jungen, Anika Witten, Monika Stoll, Isabell Schulze, Seishi Ogawa, Rainer Wiewrodt, Lara Tickenbrock, Wolfgang E. Berdel, Martin Dugas, Nils H. Thoennissen, and Carsten Müller-Tidow. Dna methyltransferase inhibition reverses epigenetically embedded phenotypes in lung cancer preferentially affecting polycomb target genes. *Clinical Cancer Research*, 20(4):814–826, 2014. ISSN 1078-0432. doi: 10.1158/1078-0432.CCR-13-1483. URL https://clincancerres.aacrjournals.org/content/20/4/814.

F. Pedregosa, G. Varoquaux, A. Gramfort, V. Michel, B. Thirion, O. Grisel, M. Blondel, P. Prettenhofer, R. Weiss, V. Dubourg, J. Vanderplas, A. Passos, D. Cournapeau, M. Brucher, M. Perrot, and E. Duchesnay. Scikit-learn: Machine learning in Python. *Journal of Machine Learning Research*, 12:2825–2830, 2011.
